# RPL26/uL24 UFMylation is essential for ribosome-associated quality control at the endoplasmic reticulum

**DOI:** 10.1101/2023.03.08.531792

**Authors:** Francesco Scavone, Samantha C. Gumbin, Paul A. DaRosa, Ron R. Kopito

## Abstract

Ribosomes that stall while translating cytosolic proteins are incapacitated by incomplete nascent chains, termed “arrest peptides” (APs) that are destroyed by the ubiquitin proteasome system (UPS) via a process known as the ribosome-associated quality control (RQC) pathway. By contrast, APs on ribosomes that stall while translocating secretory proteins into the endoplasmic reticulum (ER-APs) are shielded from cytosol by the ER membrane and the tightly sealed ribosome-translocon junction (RTJ). How this junction is breached to enable access of cytosolic UPS machinery and 26S proteasomes to translocon- and ribosome-obstructing ER-APs is not known. Here, we show that UPS and RQC-dependent degradation of ER-APs strictly requires conjugation of the ubiquitin-like (Ubl) protein UFM1 to 60S ribosomal subunits at the RTJ. Therefore, UFMylation of translocon-bound 60S subunits modulates the RTJ to promote access of proteasomes and RQC machinery to ER-APs.

**Significance Statement:** UFM1 is a ubiquitin-like protein that is selectively conjugated to the large (60S) subunit of ribosomes bound to the endoplasmic reticulum (ER), but the specific biological function of this modification is unclear. Here, we show that UFMylation facilitates proteasome-mediated degradation of arrest polypeptides (APs) which are generated following splitting of ribosomes that stall during co-translational translocation of secretory proteins into the ER. We propose that UFMylation weakens the tightly sealed ribosome-translocon junction, thereby allowing the cytosolic ubiquitin-proteasome and ribosome-associated quality control machineries to access ER-APs.

## Introduction

Life depends on the ability to produce and maintain a healthy proteome. Accordingly, all cells have protein quality control (PQC) systems that surveil the proteome, selectively destroying proteins that are unable to acquire or maintain their native three-dimensional structures because of damage or errors in synthesis, folding, or oligomeric assembly. In addition to proteome surveillance, PQC can also act on stalled nascent polypeptides at the ribosome through a process known as ribosome-associated quality control (RQC) (7–9). Ribosomes can stall during translational elongation when they encounter damaged mRNA, depleted charged aminoacyl tRNA pools, or specific mRNA sequences (10). Another common cause of ribosome stalling in eukaryotes occurs when ribosomes translate poly(A) tracts, synthesizing polylysine homopolymers. This “non-stop” translation can occur when ribosomes fail to terminate at stop codons or because of premature mRNA polyadenylation (11–14). Some poly(A) stalls can be resolved if the ribosome resumes translation in the same reading frame, i.e., “readthrough” (RT), or in an alternate, “frameshifted” (FS) reading frame. However, failure of the stalled ribosome to resume translation in a timely manner results in a collision with an upstream ribosome, creating a unique composite interface that specifically recruits ribosome rescue factors that degrade the arrested incomplete nascent peptide allowing reuse of the ribosomal subunits. Such factors include ZNF598, an E3 ubiquitin ligase (E3^Ub^) that ubiquitylates the 40S subunit of the stalled ribosome, and the ASCC/hRQT complex (ASCC3, ASCC2, TRIP4), which recognizes the ubiquitylated 40S and dissociates the stalled ribosome into 40S and 60S subunits (Fig 1A) (15–20). The released 40S subunit can be reused, but the 60S subunit remains attached to an incomplete nascent chain-peptidyl tRNA conjugate that obstructs both the P-site and the exit tunnel (ET) (21–26). This AP must be extracted from the ET and destroyed by RQC, thereby preventing toxic aggregation of the AP and allowing reuse of the 60S (27, 28).

**Figure 1:**
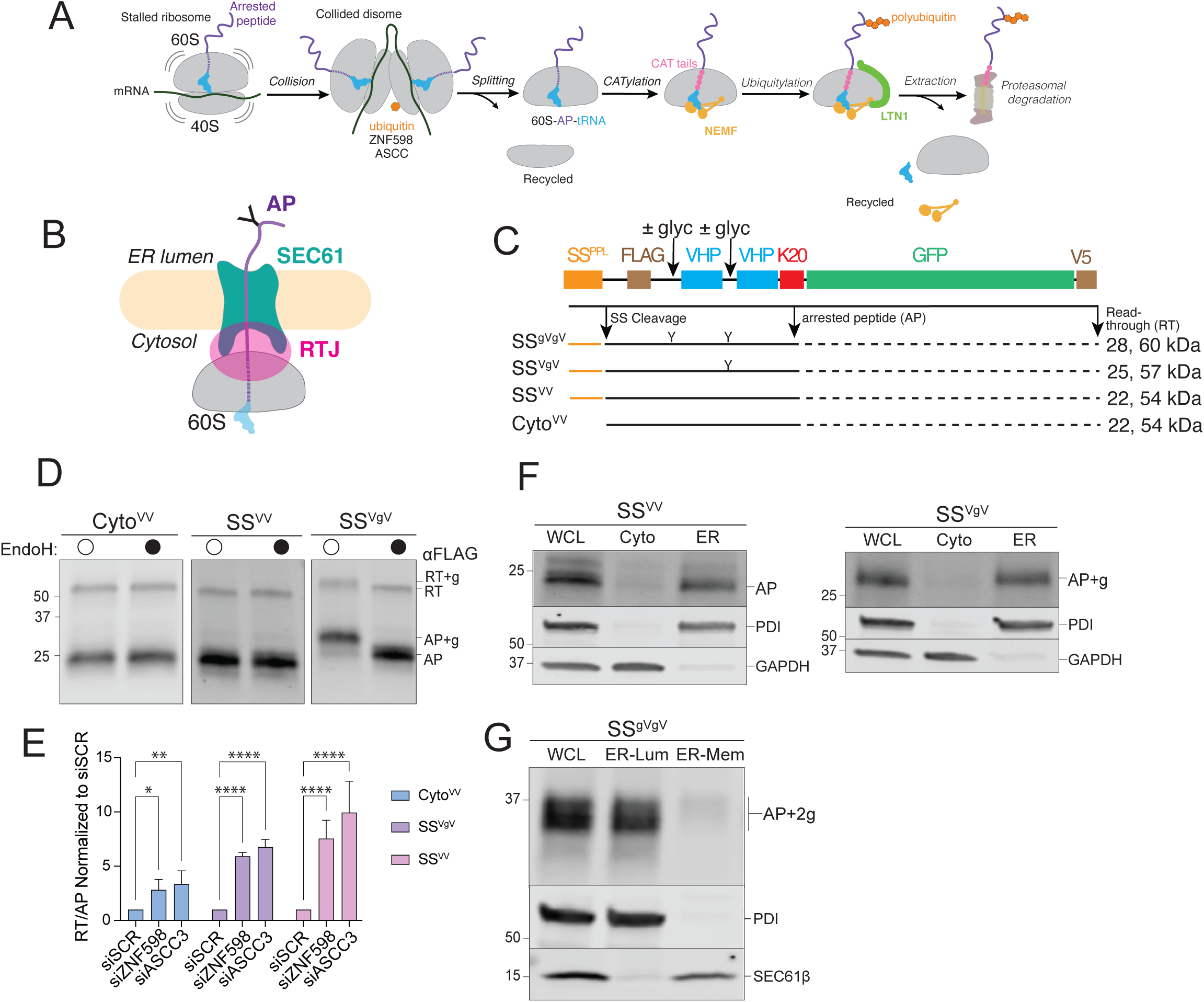
ER-targeted reporters to investigate ribosome stalling at the ER. **A:** Schematic of ribosome-associated quality control (RQC) in the mammalian cytoplasm. **B:** Topological organization of 60S-AP-tRNA stalled at an ER translocon. APs on ER ribosomes are topologically segregated from the cytosol by the ribosome-translocon junction (RTJ). **C:** Schematic of the stalling reporters used in this study. SS^PPL^, signal sequence from bovine preprolactin; FLAG, FLAG epitope tag; ± glyc, presence or absence of an N-glycosylation sequon; “Y”, N-glycan; VHP, villin headpiece domain; K20; polylysine stalling sequence of 20 lysine residues; GFP, superfolder green fluorescent protein; V5, epitope tag. Composition of each reporter shown below, with predicted MW (*indicated in kDa*) for arrest peptide (AP, black line) or readthrough (RT, black line + dashed black line) species produced by each stalling reporter. **D:** Endoglycosidase H (endoH) treatment on SS^VgV^ reporter demonstrates glycosylation of SS^VgV^ by increase in AP mobility. HEK293 cells were transfected with the indicated reporters and products were analyzed by immunoblot of whole cell lysates (WCLs) with FLAG antibody. Labels indicate mobilities of glycosylated (+g) and non-glycosylated RT and APs; data shown are representative of three independent experiments. **E:** ER-stalled ribosomes are recognized and split by ZNF598 and ASCC3, respectively. HEK293 cells were transfected with scrambled (SCR), ZNF598, or ASCC3 small interfering RNAs (siRNA) and stalling reporter constructs Cyto^VV^, SS^VV^, SS^VgV^. Quantification of RT and AP species was calculated as a ratio of RT/AP. Data are the mean ± SD of at least three independent experiments. *p < 0.05, **p < 0.01, ****p < 0.0001 determined by two-way ANOVA. **F:** Cell fractionation analysis of subcellular AP distribution shows that ER-APs co-fractionate with ER markers. U2OS cells were transfected with the indicated reporters and subjected to cell fractionation. Reporter products were analyzed by immunoblot of WCL, cytosolic (Cyto), and ER cell fractions with FLAG antibody. GAPDH: cytosol marker; PDI: ER marker; data shown are representative of three independent experiments. **G:** ER-APs predominantly localize in the ER lumen. HEK293 cells were transfected with SS^gVgV^. Cells were fractionated under conditions optimized to promote leakage of ER luminal contents while preserving membrane integrity as optimized in Fig S1E. Reporter products were analyzed by immunoblot of WCL, ER lumen (ER-Lum), and ER membrane (ER-Mem) fractions with FLAG antibody. PDI: ER lumen marker; SEC61β: ER membrane marker; data shown are representative of two independent experiments.

RQC is initiated when NEMF binds to the exposed intersubunit interface of the 60S subunit, and Listerin (LTN1), an E3^Ub^, polyubiquitylates the nascent chain, likely promoting its capture and extraction by the AAA+ ATPase p97/VCP and degradation by the 26S proteasome (21–26). NEMF and its yeast homologue Rqc2 mediate CATylation, the non-templated polymerization of amino acids to the C-terminus of the tRNA-linked AP, generating so-called CAT tails (24, 29). In mammalian cells, CAT tails are composed predominantly of alanine residues (28, 30). CATylation facilitates AP clearance in at least two ways. CAT tail formation can promote LTN1-mediated ubiquitylation of APs still bound to the 60S subunit by pushing out lysine residues, such as those encoded by poly(A), that are buried within the ET (31) or are otherwise not accessible (32). Alternatively, CAT tails can autonomously function as degrons to enable ubiquitylation by cytosolic E3s, promoting degradation of APs after their release from the 60S subunit (30, 32). How the RQC pathway resolves stalled APs on cytosolic ribosomes has been investigated in considerable detail and is the subject of several recent reviews (7–9, 33–35). By contrast, the role of RQC in resolving stalled ribosomes engaged in cotranslational translocation of proteins across the membrane of the endoplasmic reticulum (ER) has been relatively unexplored.

Ribosomes that synthesize secretory and membrane proteins at ER translocons can also stall, but how these stalls are resolved is poorly understood and complicated by the fact that ER-APs are shielded from cytosolic RQC and UPS machinery by the ribosome-translocon junction (RTJ) between the 60S and the SEC61 translocon (Fig 1B) (36–40). In yeast, ER-targeted non-stop proteins can be eliminated by a process requiring the ribosome rescue factors Dom34/Hbs1 (2) and the E3^Ub^ ligase, Listerin (Ltn1) (1). Genetic disruption of RQC in yeast impairs protein translocation into the ER, suggesting that ER-APs can obstruct translocons in addition to the ribosome ET (Fig 1B). The mechanism of ER-AP clearance in mammals is far less clear. One study concluded that RQC machinery can be recruited to SEC61 translocons in response to cycloheximide-induced global stalling, (4) potentially implicating RQC machinery in resolving ER stalls. By contrast, a more recent study reported that ER-AP degradation does not require RQC or UPS machinery but is dependent instead on lysosomes and UFMylation (5).

UFMylation (Fig S1A) refers to the process in which the small ubiquitin-like protein, UFM1, is covalently conjugated to target proteins (41, 42). UFMylation is an ancient pathway, present in the last common eukaryotic ancestor, but has been lost from three eukaryotic lineages, most notably fungi (43–45). UFM1 forms isopeptide-linked adducts with lysines on target proteins, catalyzed by dedicated E1 (UBA5), E2 (UFC1) enzymes, and a heterotrimeric E3 ligase (E3^UFM1^) composed of UFL1, DDRGK1, and CDK5RAP3 (46). UFM1 is removed from clients by a dedicated cysteine protease, UFSP2 (Fig S1A). The principal client of UFMylation is the 60S ribosomal protein, RPL26 (5, 47). UFMylation is highly selective for RPL26 lysines K132 and K134 on 60S ribosomal subunits that are docked on the ER membrane, a specificity that is conferred by the restriction of the E3^UFM1^ to the cytosolic face of the ER. Genetic disruption of UFMylation genes strongly induces ER stress, and UFM1 genes are transcriptional targets of the unfolded protein response (48–52), suggesting a functional relationship between UFMylation and ER proteostasis. Structures of ER-docked ribosomes reveal the two UFMylated RPL26 lysines to be positioned immediately adjacent to the ribosome ET and the translocon, suggesting that this modification could potentially influence the RTJ (5, 47). UFMylated RPL26 is primarily found on free 60S ribosomal subunits (47), suggesting that UFMylation plays a post-termination role. However, the function of this ribosome modification at the ER-membrane remains to be identified. Here, we sought to identify the mechanism by which ER-APs are degraded. Our data establish a central role for RQC machinery, UFMylation, and the UPS in recognizing, ubiquitylating, and degrading stalled ER-APs.

## Results

### Ribosome stalls at the ER are resolved by ribosome-associated quality control

To elucidate how nascent chains on ribosomes that stall during cotranslational translocation into the ER lumen are degraded, we engineered a set of cytosolic and ER-targeted reporters that contain a polylysine (K20) tract to mimic “non-stop” translation into poly(A). Each reporter consists of tandem 35-residue villin “headpiece” (VHP) domains that autonomously and rapidly fold into stable 3-helix bundles (53), and a downstream superfolder GFP to monitor readthrough (RT) beyond the K20 stall sequence (Fig 1C). The efficient N-terminal signal sequence (SS) from bovine preprolactin (54, 55) was appended to the N-termini to direct these reporters for cotranslational translocation into the ER. N-glycosylation sequons were included in two reporter variants, SS^VgV^ and SS^gVgV^, to monitor ER targeting and dislocation. When expressed in cells, SS^VV^ and an SS-lacking variant, Cyto^VV^, gave rise to FLAG-immunoreactive electrophoretic species with mobilities corresponding to those expected for RT and arrest peptide (AP) products (Fig 1D), while the sequon-containing variants generated AP and RT bands corresponding to the mobilities expected for singly (Fig 1D) and doubly (Fig S1B) glycosylated forms of AP and RT. To confirm the identities of these AP species, we treated cell lysates with endoglycosidases (endoH or PNGase), which cleave N-linked glycans (56, 57), causing increased mobility of the RT and AP bands (i.e., ∼2 kDa for RT+g and AP+g; ∼4 kDa for RT+2g and AP+2g), and collapsing them to the sizes observed for the corresponding non-glycosylated species for the stalling reporters (Fig 1D, S1B).

Silencing of ZNF598 or ASCC3 – which, respectively, recognize and split stalled ribosomes in the cytosol (15–20) – robustly increased the levels of both RT and frameshift (FS) species of cytosolic and ER-targeted stalling reporters, and simultaneously decreased the levels of APs (Fig 1E; S1C), confirming that ribosomes stalled during cotranslational translocation into the ER are split by the same machinery that rescues stalled ribosomes in the cytosol. Subcellular fractionation confirmed that Cyto^VV^-AP was exclusively cytosolic (Fig S1D) while the vast majority of glycosylated and non-glycosylated ER-APs co-fractionated with ER (Fig 1F). These ER-APs could either remain as translocon- and ribosome-associated nascent polypeptides at the ER membrane, or they could be released as free truncated polypeptides into the ER lumen. To distinguish between these possibilities, we titrated digitonin concentrations to identify a condition that promotes selective release of ER luminal proteins without solubilizing ER membrane proteins (Fig S1E). At 0.1% digitonin, glycosylated ER-APs robustly co-fractionated with the ER lumen protein PDI, but not with the integral membrane protein, SEC61β (Fig 1G), suggesting that glycosylated ER-APs can be released into the ER lumen. Notably, glycosylated ER-APs did not sediment with ribosomal proteins when ultracentrifuged through 1M sucrose (Fig S1F), confirming that glycosylated ER-APs were no longer bound to the 60S ribosomal subunit. Together, these data establish that ribosomes that translate ER-targeted stalling reporters are efficiently targeted to ER translocons and are split by cytosolic rescue factors to yield arrested nascent chains that can be released from the ribosome and translocon to enter the ER lumen. We conclude that these reporters are signal sequence cleaved, glycosylated, and cofractionate with the ER, making them well-suited to investigate cellular mechanisms of stall resolution for ribosomes engaged in cotranslational protein translocation into the ER.

### ER-APs are degraded by the proteasome following p97/VCP-mediated dislocation

To determine how APs on ER-stalled ribosomes are degraded, we tested the effect of bortezomib (BTZ) and bafilomycin A1 (BafA), potent and highly selective inhibitors of proteasomal and lysosomal proteolysis respectively, on the abundance of ER-targeted SS^VV^- and SS^VgV^-K20 reporters (Fig 2A). We failed to observe an effect of BafA on the levels the APs (Fig 2A), even after prolonged incubation with the drug (Fig S2A), despite a robust increase in the levels of an endogenous lysosomal substrate, LC3-II (Figs 2A, S2A). By contrast, BTZ treatment led to significant and robust increases in both Cyto^VV^- and SS^VV^-APs (Fig 2A, S2D), suggesting an essential role for proteasomes in degrading nascent chains from ribosomes that stall during cotranslational translocation into the ER. BTZ treatment on SS^VgV^ expressing cells did not significantly alter the abundance of glycosylated AP (AP+g), but instead increased the abundance of an endoH-resistant (Fig S2B) species that migrates at the molecular weight expected for signal sequence cleaved, non-glycosylated AP (Fig 2A). Because the only known cellular enzyme capable of hydrolyzing N-glycans is the cytosolic enzyme N-glycanase-1 (NGLY1) (58, 59) these data suggest that, upon stalling, some glycosylated SS^VgV^-APs are dislocated to the cytosol and deglycosylated prior to degradation by the proteasome. To confirm this, we used subcellular fractionation to assess the effect of BTZ on AP partitioning between ER and cytosol. As expected, Cyto^VV^-APs were stabilized in the cytosol following BTZ treatment (Fig S2C). In untreated cells, APs from ER-targeted reporters were exclusively ER-associated and, for SS^VgV^, fully glycosylated (Fig 2B; see also Fig 1D, F). However, the BTZ-stabilized APs of both ER-targeted reporters robustly co-fractionated with cytosolic markers (Fig 2B), confirming that ER-stalled APs are degraded by the proteasome following dislocation to the cytosol. Treatment with the p97/VCP AAA+ ATPase inhibitor NMS-873 substantially reduced the BTZ-promoted accumulation of deglycosylated SS^VgV^-AP in the cytosol (Fig 2C), indicating that p97/VCP is required to dislocate glycosylated APs across the ER membrane. These data establish that ER-APs are degraded by a mechanism that requires the UPS, a conclusion that differs sharply from that reached in a previous study (5), which reported that APs derived from an ER-targeted stalling reporter, ER-K20 are degraded in lysosomes. The most notable difference between ER-K20 and the reporters used in our study is the presence of a luminal GFP moiety in ER-K20. We considered that the intrinsic stability of the GFP β-barrel might make ER-K20 less susceptible to cytosolic dislocation, causing it to be preferentially released into the ER lumen and hence, escape proteasomal degradation. To test this, we replaced the VHP domains in SS^VV^ with GFP. This reporter, SS^GFP^, was unaffected by BafA, but robustly stabilized by BTZ treatment (Fig S2E). Therefore, the presence of structurally diverse, tightly folded domains in the luminal portion of the ER-AP does not affect the dependence on the proteasome for degradation.

**Figure 2:**
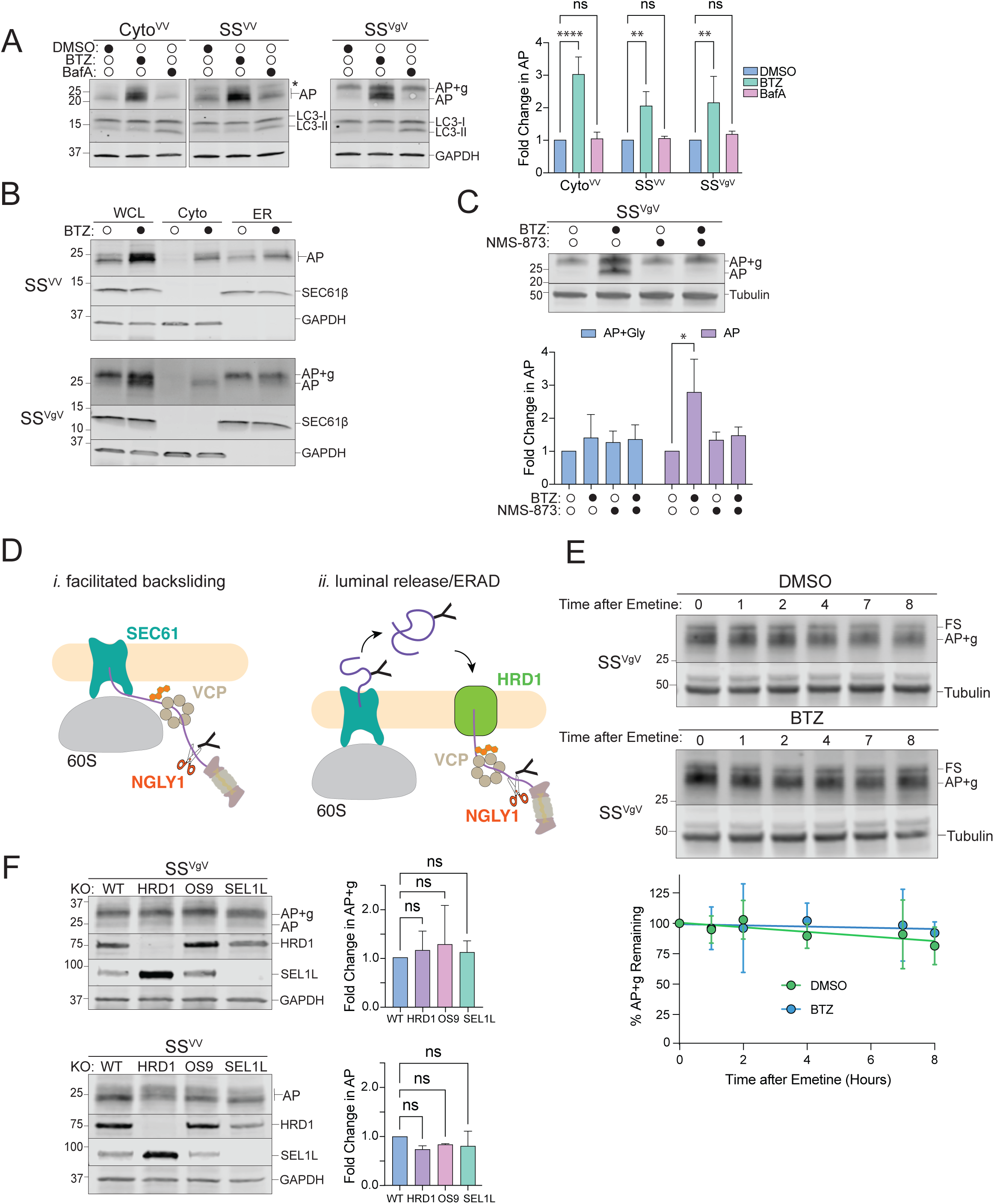
Proteasomal degradation of ER-AP requires dislocation to the cytosol via a p97/VCP-dependent and HRD1-independent pathway. **A:** Cytosolic- and ER-APs are stabilized by proteasome but not lysosome inhibitors. *Left panels,* HEK293 cells expressing the indicated reporters were treated either with DMSO, 1 µM bortezomib (BTZ), or 100 nM Bafilomycin A1 (BafA) for 4 hr. WCLs were analyzed by immunoblot with FLAG antibody to detect AP and AP+g species, and LC3 antibody to assess the effect of BafA treatment on LC3-II accumulation. Asterisks indicate a nonspecific immunoreactive band. GAPDH: loading control. *Right panel,* Quantification of immunoblot data as indicated. Fold change in AP relative to DMSO treated cells was calculated after normalization to GAPDH. Data are the mean ± SD of at least three independent experiments. **p < 0.01, ****p < 0.0001 determined by two-way ANOVA. **B:** ER-APs are dislocated to the cytosol prior to degradation via the proteasome. U2OS cells were transfected with the indicated reporter and treated with DMSO or 1 µM BTZ for 4 hr prior to cell fractionation. Reporter products were analyzed by immunoblot of WCL, Cyto, and ER cell fractions with FLAG antibody. GAPDH: cytosol marker; SEC61β: ER marker; data shown are representative of three independent experiments. **C:** ER-AP dislocation to the cytosol depends on p97/VCP. *Upper panel,* HEK293 cells expressing SS^VgV^ were treated with 1 µM BTZ, 5 µM NMS-873, or 1 µM BTZ and 5 µM NMS-873 for 4 hr. Reporter products were analyzed by immunoblot with FLAG antibody. Tubulin: loading control. *Lower panel,* Quantification of immunoblot data as indicated. Data are the mean ± SD of two independent experiments. *p < 0.05 determined by two-way ANOVA. **D:** Two models of ER-AP dislocation to the cytosol via p97/VCP and degradation by the proteasome. Details in text. **E:** Turnover of glycosylated SS^VgV^ is slow. *Upper panels,* HEK293 cells transfected with SS^VgV^ (“Original” reporter, see figure S2F) were treated with 20 µM emetine and either DMSO or 1 µM BTZ for the indicated times. Reporter products were analyzed by immunoblot with FLAG antibody. Tubulin: loading control. *Lower panel,* Quantification of immunoblot data as indicated. %AP+g remaining was determined by normalizing AP+g signal to tubulin signal, then calculating the fraction remaining relative to time = 0 hr. Data are the mean ± SD of two independent experiments. **F:** ER-AP degradation does not require the HRD1 retrotranslocon. *Left panels,* HEK293 WT, *HRD1^KO^*, *OS9^KO^*, and *SEL1L^KO^* cell lines were transfected with the indicated reporters. Reporter products were analyzed by immunoblot with FLAG antibody. Knockouts were confirmed by blotting with antibodies against endogenous HRD1 or SEL1L proteins. GAPDH: loading control. *Right panels,* Quantification of AP intensity for SS^VV^ and AP+g intensity for SS^VgV^. Fold changes relative to WT cells were calculated after normalization to GAPDH. Data are the mean ± SD of at least two independent experiments. ns > 0.05, determined by one-way ANOVA. **F:** Schematic of the two reporter variants illustrating the frameshift (FS) species generated by our stalling reporters as described in the materials and methods section. Original: frameshift product generated by this reporter is ∼25kD. FS-corrected: frameshift product generated by this reporter is 60kD or 65kD. S Tag+1 and S Tag+2 are generated by out of frame translation downstream of the stalling sequence (K20). The Original reporter was used in Figures 2E, S2A. The FS-corrected reporter is used in all other experiments.

The most direct route by which APs that stall on translocon-docked 60S ribosomes could access the cytosol is via ATP-dependent, p97/VCP-facilitated backsliding through SEC61 (Fig 2D, model *i, facilitated backsliding*). Alternatively, these ER-APs could be first released into the lumen and then dislocated to the cytosol by the HRD1 retrotranslocon (Fig 2D, model *ii, luminal release/ERAD*). The requirement for p97/VCP (Fig 2C) does not distinguish between these two alternatives as both RQC (25, 26) and ERAD (60–63) exploit this ubiquitin-dependent segregase/unfoldase activity. Translation shut-off experiments revealed luminal AP+g to be stable (t_1/2_ > 8hr) and negligibly affected by BTZ treatment, indicating that, once released into the ER-lumen, it is not secreted, degraded by the lysosome, or retrotranslocated by ERAD (Fig 2E). This conclusion is supported by the observation that disruption of genes encoding HRD1, OS9, or SEL1L, essential components of the HRD1 ERAD retrotranslocon complex (36, 64), did not significantly elevate either SS^VV^- or SS^VgV^-AP levels (Fig 2F), confirming that ERAD does not contribute substantially to degradation of these APs. The stability of the luminally released AP is not surprising because our reporters are composed entirely of folded helical domains interspersed with short unstructured linkers (Fig 1C) and thus lack ERAD-promoting degrons like exposed hydrophobic patches or interrupted secondary structure elements (65). Together, these data strongly suggest a model in which translocon-engaged APs can partition between two fates: release into the ER lumen, where ER-APs accumulate in the absence of a degron – or extraction from the SEC61 translocon into the cytosol by p97/VCP to be rapidly degraded by the proteasome.

### ER-AP degradation requires cytosolic RQC machinery

Since p97/VCP recognizes clients via ubiquitin chains (66), these data suggest a role for a ubiquitin ligase distinct from HRD1 in ER-AP clearance. Degradation of non-stop membrane and secretory proteins in yeast is also independent of Hrd1 but requires the RQC E3^Ub^, Ltn1 (1, 3). Although a direct role for the mammalian ortholog, Listerin (LTN1), in ER-RQC has not been established, a previous study found that the RQC components NEMF and LTN1 are recruited to ER membranes in response to translational stalling induced by sub-stoichiometric concentrations of cycloheximide (4). While HRD1 silencing failed to stabilize either cytosolic or ER-APs, we observed that LTN1 disruption led to significant elevation of cytosolic-(Cyto^VV^) and ER-APs (SS^VV^ and SS^VgV^) (Fig 3A, S3A). We also observed elevated levels of cytosolic-(Cyto^VV^) and ER-APs (SS^VV^ and SS^VgV^) in clonal lines harboring CRISPR/Cas9-directed knockout of *LTN1* (Fig 3B) and *NEMF* (Fig 3C). Thus, both LTN1 and NEMF are required to degrade APs stalled on ribosomes engaged in cotranslational translocation of secretory proteins at the ER. Importantly, the ER-APs that were stabilized in *NEMF* and *LTN1* knockouts accumulated predominantly in the ER fraction (Fig 3D), suggesting that failure of RQC leads to increased release of APs into the ER lumen. These data support the conclusion that NEMF and LTN1 are required for dislocation of ER-stalled APs to the cytosol.

**Figure 3:**
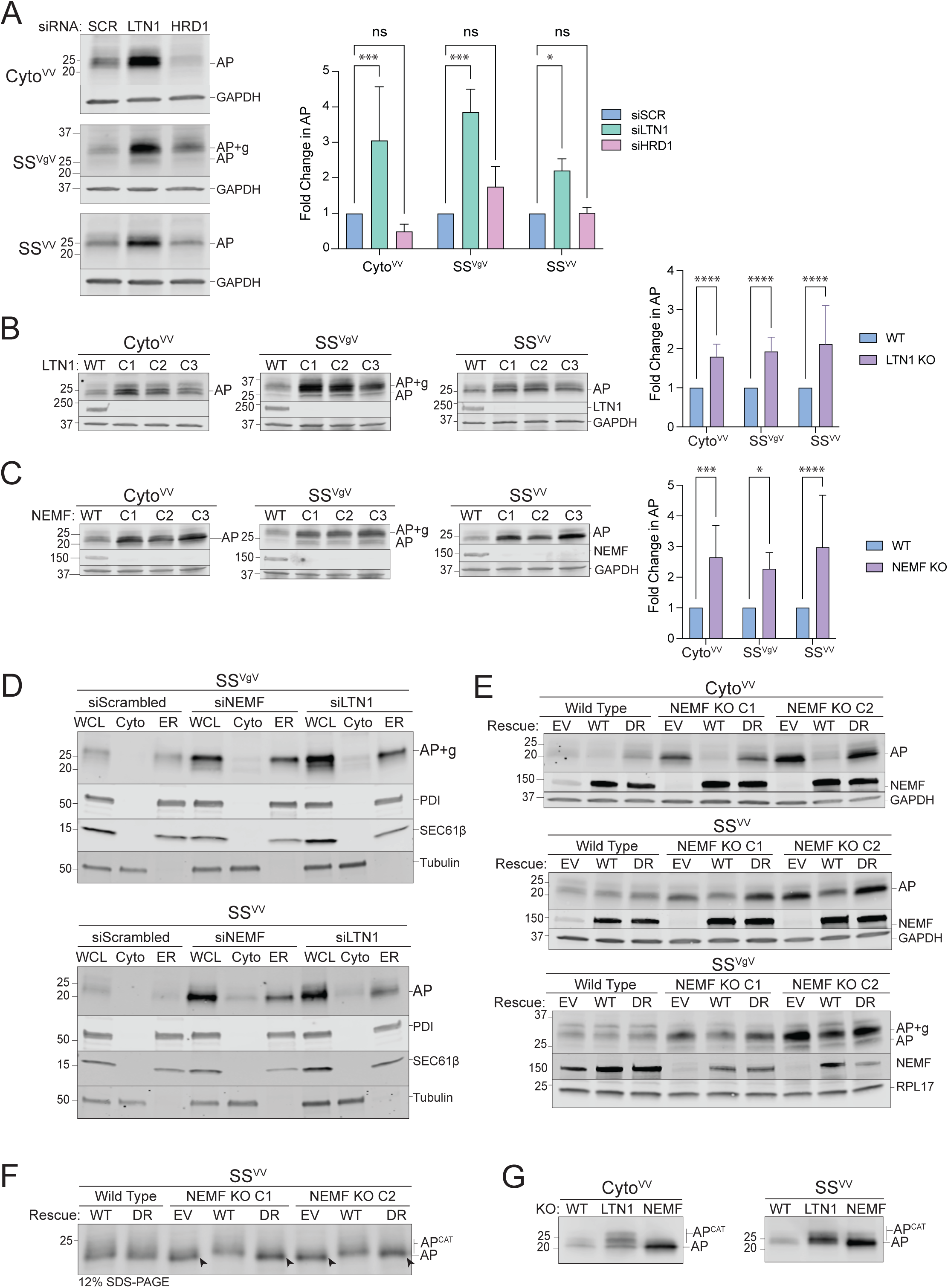
ER-AP degradation requires RQC machinery. **A:** Knockdown of LTN1 but not HRD1 stabilizes both cytosolic- and ER-APs. *Left panels,* HEK293 cells were transfected with the indicated siRNAs and the indicated stalling reporters. Reporter products were analyzed by immunoblot with FLAG antibody. GAPDH: loading control. *Right panel,* Quantification of AP intensity for Cyto^VV^ and SS^VV^ and AP+g intensity for SS^VgV^. Fold change relative to WT cells was calculated after normalization to GAPDH. Data are the mean ± SD of at least 3 independent experiments. *p < 0.05, ***p < 0.001, determined by two-way ANOVA. **B:** LTN1 is required for degradation of cytosolic- and ER-APs. *Left panels,* HEK293 WT and clonal *LTN1^KO^* cell lines (C1, C2, C3) were transfected with the indicated reporters. Reporter products were analyzed by immunoblot with FLAG antibody. Knockouts were confirmed by blotting with antibodies against endogenous LTN1 protein. GAPDH: loading control. *Right panel,* Quantification as in Fig 3A. Fold change relative to WT cells was calculated after normalization to GAPDH. Data are the mean ± SD of at least 3 independent experiments. ****p < 0.0001, determined by two-way ANOVA. **C:** NEMF is required for degradation of cytosolic- and ER-APs. *Left panels,* HEK293 WT and clonal *NEMF^KO^* cell lines (C1, C2, C3) were transfected with the indicated reporters. Reporter products were analyzed by immunoblot with FLAG antibody. Knockouts were confirmed by blotting with antibodies against endogenous NEMF protein. GAPDH: loading control. *Right panel,* Quantification as in Fig 3A. Fold change relative to WT cells was calculated after normalization to GAPDH. Data are the mean ± SD of at least 3 independent experiments. *p < 0.05, ***p < 0.001, ****p < 0.0001, determined by two-way ANOVA. **D:** Disruption of RQC favors luminal release of ER-APs. U2OS cells were transfected with indicated siRNAs and stalling reporters followed by cell fractionation. Reporter products were analyzed by immunoblot of WCL, Cyto, and ER cell fractions with FLAG antibody. Tubulin: cytosol marker; SEC61β and PDI: ER markers. **E:** CATylation is required for ER-AP degradation. HEK293 WT or clonal *NEMF^KO^* cells (C1 and C2) were transfected with empty vector (EV), WT NEMF, or NEMF-DR and the indicated stalling reporters. Reporter products were analyzed by immunoblot with FLAG antibody. Endogenous and ectopic NEMF expression validated by immunoblotting with anti-NEMF antibody. GAPDH or RPL17: loading controls. **F:** ER-APs are CATylated. HEK293 WT or clonal *NEMF^KO^* cells (C1 and C2) were transfected with EV, WT, or DR NEMF and the indicated stalling reporters as in panel E. SS^VV^-transfected cell lysates were separated by 12% SDS-PAGE and analyzed by immunoblot with FLAG antibody. Unmodified APs are indicated by the label “AP” and by arrowheads; CATylated APs are indicated by “AP^CAT^”; data shown are representative of two independent experiments. **G:** CATylated ER-APs accumulate in LTN1-deficient cells. HEK293 cells were transfected with the indicated siRNAs and stalling reporters. Reporter products were analyzed by immunoblot with FLAG antibody. AP and AP^CAT^ labels indicate unmodified and CATylated APs, respectively; data shown are representative of two independent experiments.

NEMF specifically binds to 60S-peptidyl tRNA complexes at the P-site, thereby discriminating between empty 60S subunits and those with stalled nascent chains, and facilitates recruitment of LTN1 (21, 25). NEMF also elongates APs by polymerizing the addition of CAT tails to the C-termini of stalled APs (28, 30). To determine whether NEMF also CATylates ER-APs, we compared AP levels in *NEMF^KO^* cells that were rescued with either WT NEMF or with a CATylation defective variant (D96A/R97A; “DR”) (28, 30) (Fig 3E). As expected, the elevated abundance of Cyto^VV^-AP in two independent clonal *NEMF^KO^* lines was fully rescued by expression of WT NEMF but not NEMF-DR. Similarly, WT NEMF, but not NEMF-DR, rescued the elevated levels of SS^VV^-AP or SS^VgV^-AP in two independent clonal *NEMF^KO^* lines (Fig 3E). These data suggest an essential role for CATylation in resolving stalls at the ER. As an independent confirmation, we observed a ∼1.3 kDa decrease in mobility of SS^VV^-AP in *NEMF^KO^* cells rescued with WT NEMF but not with the DR variant (Fig 3F, S3B-D). Moreover, the mobilities of both cytosolic- and ER-APs were further decreased in *LTN1^KO^* relative to WT cells (Fig 3G), consistent with LTN1 functioning downstream of NEMF to promote degradation of CATylated APs. Together, these data strongly support the conclusion that both NEMF and LTN1 are required to facilitate ubiquitin-dependent cytoplasmic extraction of arrested nascent chains from stalled ribosomes docked at ER translocons.

### Ribosome stalling at the ER promotes RPL26 UFMylation

The finding that RQC machinery is required to dislocate APs from ribosomes stalled at the ER begs the question of how p97/VCP and the UPS can access nascent chains that are shielded from the cytosol by the RTJ (Fig 1B). A possible role for UFMylation in this process was previously suggested by the findings that UFMylation is stimulated by ribosome collisions at the ER (5), and that UFM1 is conjugated to a single ribosomal protein, RPL26, at the RTJ (5, 47). To investigate the relationship between UFMylation and RQC we exposed cells to anisomycin (ANS), an elongation inhibitor that, at sub-stoichiometric concentrations, stochastically stalls a subset of ribosomes causing non-inhibited upstream ribosomes to collide with them, triggering RQC (18, 67). Brief (20 min) exposure to ANS stimulated a dose-dependent increase in UFMylated RPL26 that was maximal at 200 nM, a concentration previously reported to be optimal to induce collisions on cytosolic ribosomes (Fig 4A). Although ANS treatment stalls ribosomes globally, elevated RPL26 UFMylation was observed only on ER-associated ribosomes (Fig 4B), consistent with localization of the E3^UFM1^ to this organelle. This suggests that UFMylation is stimulated in response to stalling of ribosomes engaged in cotranslational translocation across the ER membrane.

**Figure 4:**
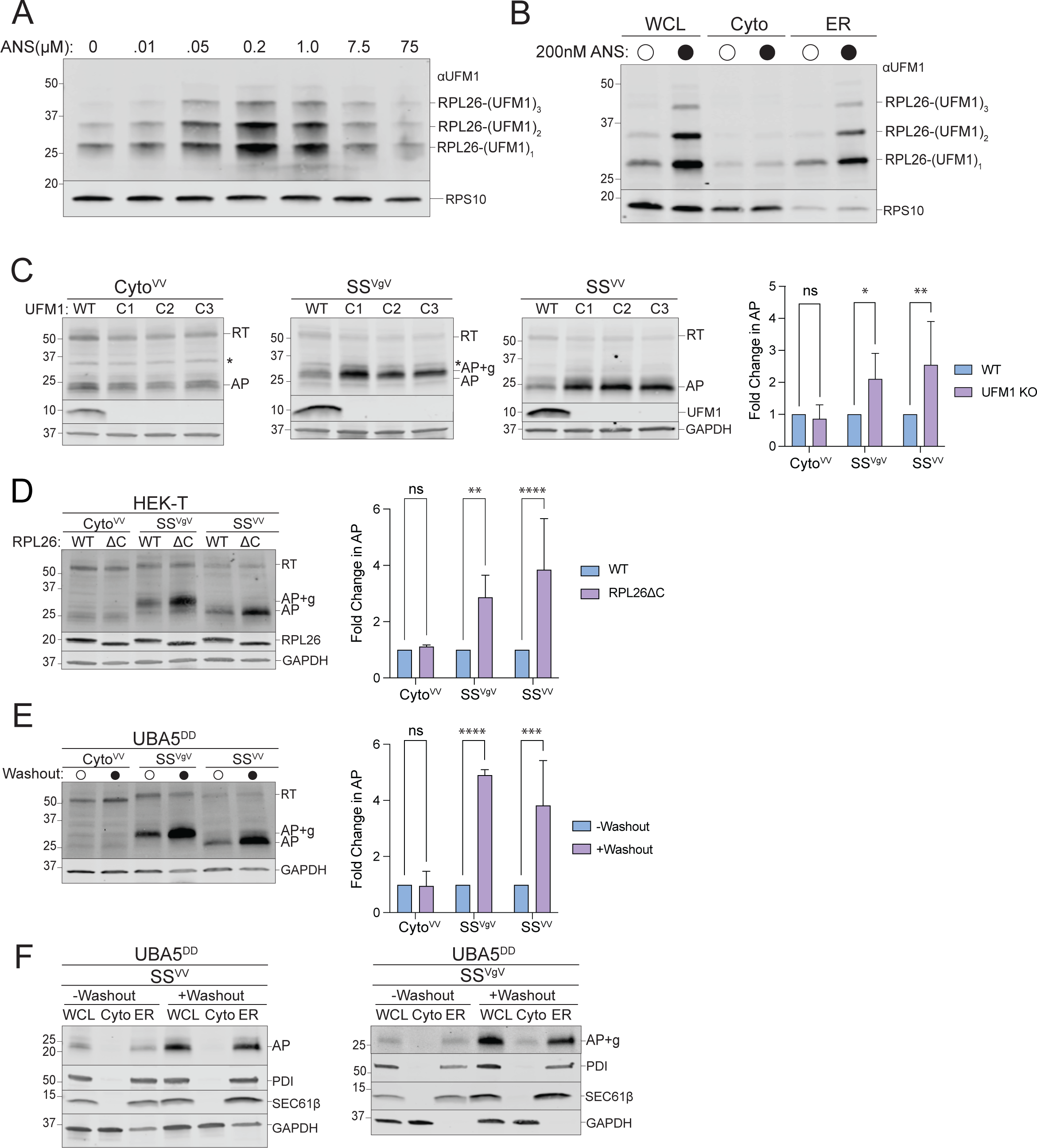
ER-AP degradation requires UFMylation. **A:** Global ribosome stalling promotes RPL26 UFMylation. U2OS cells were treated with the indicated concentrations of anisomycin (ANS) for 20 min treatment prior to harvesting. Cell lysates were sedimented through a sucrose cushion and the pellets were analyzed by immunoblot using an anti-UFM1 antibody. RPS10: loading control. **B:** Stalling induced RPL26 UFMylation occurs primarily on ER-bound ribosomes. U2OS cells were treated with either DMSO or 200 nM ANS for 20 min prior to cell fractionation and immunoblotting with UFM1. RPS10: loading control; data shown are representative of two independent experiments. **C:** Knockout of *UFM1* stabilizes ER-but not cytosolic-APs. *Left panels,* HEK293 WT and clonal *UFM1^KO^* cell lines (C1, C2, C3) were transfected with the indicated reporters. Reporter products were analyzed by immunoblot with FLAG antibody. Asterisks indicate a nonspecific immunoreactive band. Knockouts were confirmed by blotting with antibodies against endogenous UFM1 protein. GAPDH: loading control. *Right panel,* Quantification of AP intensity for Cyto^VV^ and SS^VV^ and AP+g intensity for SS^VgV^. Fold change relative to WT cells was calculated after normalization to GAPDH. Data are the mean ± SD of at least 3 independent experiments. ns > 0.05, *p < 0.05, **p < 0.01, determined by two-way ANOVA. **D**: Specific UFMylation of RPL26 is required for ER-AP degradation. *Left panel,* HEK-T WT and clonal RPL26ΔC cell lines were transfected with the indicated reporters. Reporter products were analyzed by immunoblot with FLAG antibody. C-terminal deletion of RPL26 was confirmed by blotting with an antibody against endogenous RPL26 protein. GAPDH: loading control. *Right panel,* Quantification of AP intensity for Cyto^VV^ and SS^VV^ and AP+g intensity for SS^VgV^. Fold change relative to WT cells was calculated after normalization to GAPDH. Data are the mean ± SD of at least 3 independent experiments. ns > 0.05, **p < 0.01, ****p < 0.0001 determined by two-way ANOVA. **E:** Acute disruption of UFMylation stabilizes ER-but not cytosolic-APs. *Left panel,* U2OS UBA5^DD^ cells were cultured in complete DMEM with TMP (“-Washout”) or washed to remove TMP from the media (“+Washout”) before transfection with the indicated reporters. Reporter products were analyzed by immunoblot with FLAG antibody. GAPDH: loading control. *Right panel,* Quantification as in Fig 4C. Fold change relative to WT cells was calculated after normalization to GAPDH. Data are the mean ± SD of at least 3 independent experiments. ns > 0.05, ***p < 0.001, ****p < 0.0001, determined by two-way ANOVA. **F:** Acute disruption of UFMylation favors release of ER-APs into ER lumen. As in Fig 4D, followed by cell fractionation. Reporter products were analyzed by immunoblot of WCL, Cyto, and ER cell fractions with FLAG antibody. GAPDH: cytosol marker; SEC61β and PDI: ER markers; data shown are representative of two independent experiments.

To test this hypothesis, we asked whether expression of our ER-targeted stalling reporters also leads to increased RPL26 UFMylation. Expression of ER-targeted stalling reporters (SS^VV^-K20 and SS^VgV^-K20), but not cytosolic (Cyto^VV^-K20) or non-stall (K0) versions, led to substantial increases in RPL26 UFMylation (Fig S4A) that was comparable in magnitude to that observed in response to ANS treatment. RPL26 UFMylation was unaffected by cytosolic stalling, even when Cyto^VV^-K20 is strongly overexpressed (Fig S4B). In addition, RPL26 UFMylation was stimulated by expression of other ER-targeted polylysine-based stalling constructs including a stalling version of a natively secreted protein, preprolactin (PPL-K20), and an engineered type II transmembrane reporter, TM^VgV^-K20 (Fig S4C). These data confirm that collisions of ribosomes engaged in cotranslational translocation of diverse proteins into or across the ER membrane robustly and specifically induces RPL26 UFMylation.

### RPL26 UFMylation is required for ER-AP degradation

Because enhanced RPL26 UFMylation is a specific response to stalling of ribosomes engaged in synthesis of proteins at ER translocons, we sought to determine whether RPL26 UFMylation influences the stability of ER-APs. Levels of ER-, but not cytosolic-APs, were significantly elevated when the reporters were expressed in HEK293 (Fig 4C) or U2OS (Fig S4D) *UFM1^KO^* cells. ER-APs were also specifically stabilized in cells harboring stable knockouts of the UFM1 E2 and E3, *UFC1* and *UFL1*, respectively (Fig S4E). Importantly, this impairment of ER-RQC is dependent on UFM1 conjugation to RPL26, as cells lacking the sites of RPL26 UFMylation (RPL26ΔC) (5) also accumulated ER-, but not cytosolic-APs (Fig 4D).

Genes encoding UFM1 or its conjugation machinery are designated on DepMap as “common essential” (68, 69) and cell lines harboring stable knockouts of UFMylation pathway genes, while viable, exhibit altered gene expression (47). These changes include a host of ER-related phenotypes including elevated ER stress markers and impaired ERAD, likely the consequence of compensatory adaptive changes. To ensure that the observed elevation in steady-state ER-AP levels is a direct consequence of loss of UFM1 conjugation, not a secondary adaptive effect, we generated UBA5^DD^ cell lines in which the endogenous *UBA5* gene, which encodes the UFM1 E1 enzyme (Fig S1A), was replaced with a variant *UBA5* genetically fused to the *E. coli* dihydrofolate reductase (DHFR) “destabilizing domain” (DD) that causes the fusion protein to be acutely degraded following washout of the stabilizing ligand, tetramethylpyrazine (TMP) (70). Importantly, the genetic replacement strategy ensured that these cells were never exposed to even transient *UBA5* disruption during creation of the line and hence are unlikely to harbor adaptive changes due to disrupted UFMylation. Control experiments confirmed that UBA5^DD^ was > 95% eliminated within 4 hr following TMP washout (Fig S4F). The elimination of detectable UBA5^DD^ protein was accompanied by gradual loss of UFM1 conjugates to RPL26, UFC1, and UBA5, with complete loss by 16 hr (Fig S4F). Strikingly, we found that acute disruption of UBA5 resulted in increased steady-state levels of ER-APs, but not cytosolic-APs, to levels comparable to those observed in cells harboring stable disruption of *UFM1* or UFM1 conjugation machinery (Fig 4E, S4E). Importantly, the ER-APs that were stabilized following acute UBA5 disruption were enriched specifically in the ER fraction (Fig 4F), suggesting that UFM1, like NEMF and LTN1, is required to facilitate cytosolic backsliding of ER-APs. Together, these data establish that UFM1 conjugation to RPL26 is specifically required for RQC-mediated degradation of ER-, but not cytosolic-APs.

### UFMylation collaborates with RQC to degrade ER-stalled APs

The preceding data establish that RQC machinery and UFMylation are required to degrade APs on translocon-stalled ribosomes but do not address how these two systems act in concert with one another. We found that neither basal nor ANS-stimulated RPL26 UFMylation was affected by NEMF knockdown (Fig 5A), establishing that UFM1 conjugation occurs independently of NEMF function. Moreover, SS^VV^-AP was CATylated in *UFM1^KO^* cells to an extent indistinguishable from that observed in *LTN1*^KO^ cells (Fig 5B), demonstrating that RPL26 UFMylation is not required for CATylation of ER-APs, but is required for degradation of CATylated ER-APs. Although RPL26 UFMylation occurs independently of NEMF function, both UFM1 and LTN1 act downstream of NEMF-mediated CATylation to promote AP degradation. To investigate this relationship more closely we sought to determine whether simultaneous disruption of *UFM1* together with *LTN1* or *NEMF* stabilized ER-APs more than disrupting either gene alone. Unfortunately, because *NEMF^KO^* cells have a growth defect (71), which is further exacerbated by additional disruption of UFMylation, it was not possible to test the epistatic relationship between *NEMF* and *UFM1*. Silencing LTN1 in *UFM1^KO^* cells, however, did not compromise viability and exhibited no further stabilization of SS^VgV^- or SS^VV^-APs compared to the effect of disruption of either gene alone (Fig 5C). Likewise, acute inactivation of UFMylation by loss of UBA5^DD^ stabilized both ER-APs to the same extent in control or LTN1 silenced cells with TMP washout (Fig 5D). Together, these data indicate that UFMylation is required for degradation of APs stalled at ER translocons via an RQC- and UPS-dependent mechanism. UFM1 modifies the 60S at the RTJ (5, 47), raising the possibility that this modification could, either by itself or by recruiting an UFM1-interacting protein (“UFIP” in Fig 5E, *iii*) weaken the interaction between the ribosomal subunit and the translocon thus allowing p97/VCP and the UPS to access the previously occluded ER-AP (Fig 5E).

**Figure 5:**
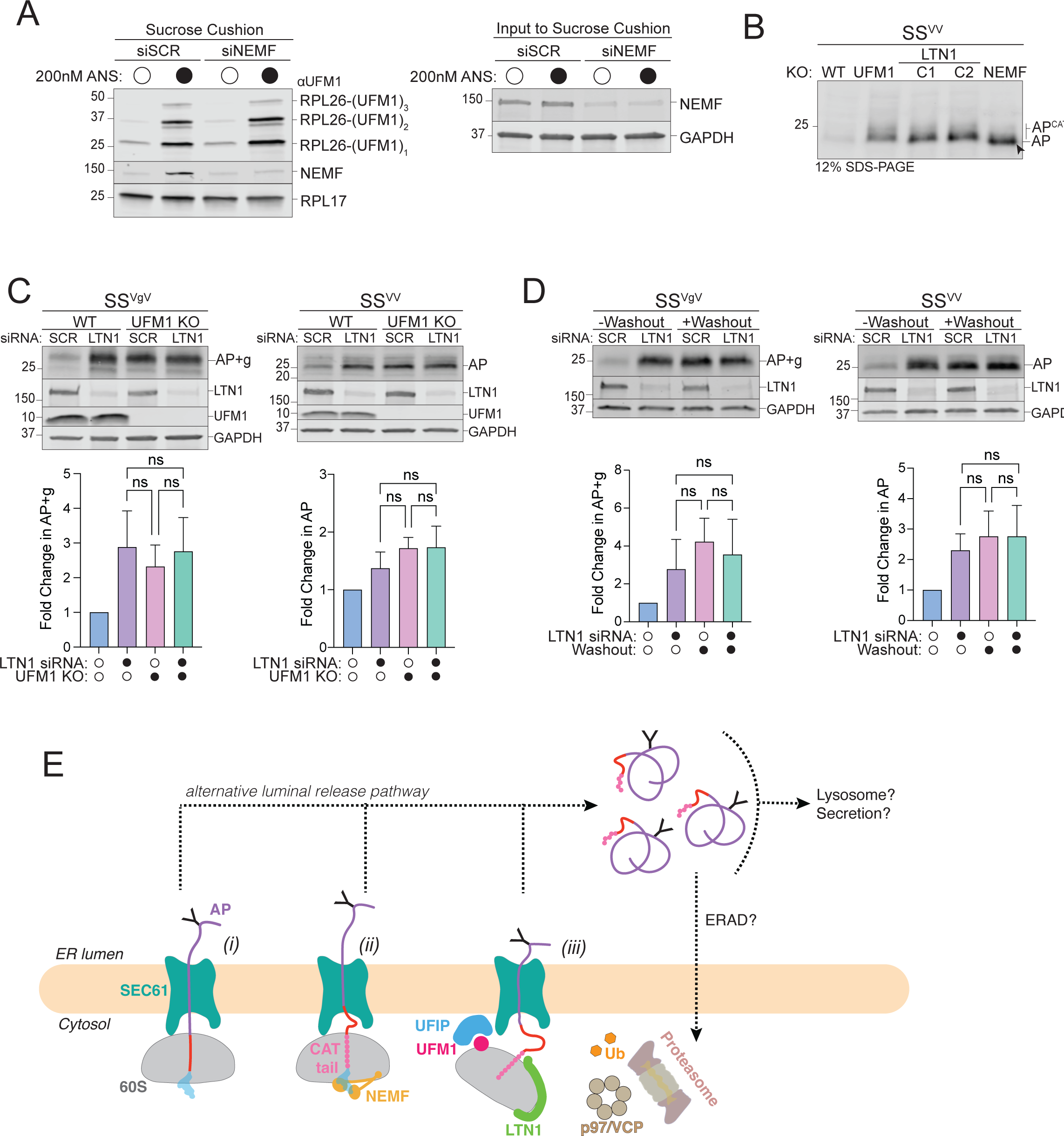
RQC machinery cooperates with UFMylation to degrade ER-APs. **A:** NEMF is not required for RPL26 UFMylation. *Left panel,* Sucrose cushion sedimentation of WCLs derived from HEK293 cells transfected with either scrambled (SCR) or NEMF siRNA and treated with either DMSO or 200 nM ANS for 30 min to induce RPL26 UFMylation. Pellets were immunoblotted with anti-UFM1 and anti-NEMF antibodies. RPL17: loading control. *Right panel,* Effect of scrambled (SCR) or NEMF siRNA on endogenous protein levels, assayed by immunoblot for endogenous NEMF with anti-NEMF antibody in WCLs (input to sucrose cushion). GAPDH: loading control. **B:** ER-AP CATylation is independent of UFMylation. HEK293 WT, *NEMF^KO^*, *LTN1^KO^* (two independent clonal lines, C1 and C2), and *UFM1^KO^* cells were transfected with SS^VV^. WCLs were separated on 12% SDS-PAGE and analyzed by immunoblot with anti-FLAG antibody. Unmodified AP are indicated by the label “AP” and by arrowheads; CATylated AP are indicated by “AP^CAT^”; data shown are representative of two independent experiments. **C:** UFM1 and LTN1 act in the same pathway to degrade ER-APs. *Upper panels,* WT or *UFM1^KO^* HEK293 cells were transfected with either scrambled (SCR) or LTN1 siRNA and the indicated reporters. Reporter products were analyzed by immunoblot with FLAG antibody. Knockout or knockdown was confirmed by immunoblot for endogenous LTN1 or UFM1 proteins with anti-LTN1 or anti-UFM1 antibodies, respectively. GAPDH: loading control. *Lower panels,* Quantification of AP intensity for SS^VV^ and AP+g intensity for SS^VgV^. Fold change relative to WT cells was calculated after normalization to GAPDH. Data are the mean ± SD of at least 2 independent experiments. ns > 0.05, determined by one-way ANOVA. **D:** UBA5 and LTN1 act in the same pathway to degrade ER-APs. *Upper panels,* U2OS UBA5^DD^ cells were transfected with either scrambled (SCR) or LTN1 siRNA and the indicated reporters. Cells were cultured in complete DMEM with TMP (“-Washout”) or washed to remove TMP from the media (“+Washout”). Reporter products were analyzed by immunoblot with FLAG antibody. Knockdown was confirmed by immunoblot for the endogenous LTN1 protein with anti-LTN1 antibody. GAPDH: loading control. *Lower panels,* Quantification as in Fig 5C. Fold change for (“+Washout”) relative to control (“-Washout”) was calculated after normalization to GAPDH. Data are the mean ± SD of at least two independent experiments. ns > 0.05, determined by one-way ANOVA. **E:** Model of ER-AP degradation. ER-APs partition between facilitated backsliding through SEC61 to cytosol (*steps i-iii*) or release into the ER lumen (*dashed lines*). Once released into the ER lumen, ER-APs are stable but could be subject to trafficking or other degradation pathways. UFIP: UFM1 interacting protein. See text for details.

## Discussion

Ribosomes that stall during protein synthesis are incapacitated by toxic tRNA-bound APs which must be destroyed in order to avert proteotoxicity and to enable reuse of 60S subunits (7, 9, 72). When stalls occur during translation of cytosolic proteins, these APs are efficiently recognized and ubiquitylated by the well-defined RQC pathway and destroyed by the 26S proteasome, releasing a “rescued” 60S ribosomal subunit competent to undergo further rounds of protein synthesis (7–10, 33–35) (Fig 1A). However, stalls that occur on ribosomes engaged in cotranslational translocation of proteins into the ER generate ER-APs that are shielded from cytosolic UPS machinery by the RTJ (Fig 1B) and disrupt protein translocation in addition to ribosome function (1, 2). While limited data in yeast suggests that some ER-APs can be cleared by a Ltn1-dependent process (1–3), how such stalls are resolved in mammalian cells has not been systematically investigated, and the few published reports reach opposing conclusions (4–6). The data reported here unambiguously establish an essential role for the RQC components NEMF and LTN1, p97/VCP, and the 26S proteasome in degrading ER-APs in mammalian cells. Moreover, we uncover a central role for UFMylation in facilitating this degradative process.

The findings presented here are integrated into a framework (Fig 5E) which schematically depicts the fates of ER-stalled APs and reconciles our findings with those from previous studies. Collided ribosomes at the ER, split by ZNF598 and ASCC3, yield a 60S-AP-tRNA adduct with the partially translocated nascent polypeptide initially extending from the P-site through the ET and translocon into the ER lumen (Fig 5E-*stage i*), a topology validated in our experiments by subcellular fractionation and glycosylation analysis (Fig 1, S1). This ER-associated ribosome stalled nascent chain is recognized by NEMF, which mediates CATylation of the ER-AP (*stage ii*) to facilitate LTN1-catalyzed ubiquitylation and proteasomal degradation (*stage iii*). In yeast, loss of Ltn1-mediated dislocation of ER-RQC clients results in release of non-stop APs into the ER lumen (1). We also find that a fraction of ER-APs partition spontaneously into the lumen (Fig 1F, G) and this fraction is substantially increased in cells with impaired RQC or UFMylation machinery (Fig 3D, 4F). Thus, ER-APs partition between (1) backsliding to the cytosol via a UFM1-dependent process requiring RQC machinery and force generation through ubiquitin-dependent ATPases such as p97/VCP or the 26S proteasome (Fig 5E, stage *iii*) and (2) release into the ER lumen (Fig 5E, dashed lines). Partitioning between these fates is likely to be determined by the relative kinetics of AP release from the 60S following hydrolysis of the peptidyl tRNA linkage, ER-AP ubiquitylation, and extraction (Fig 5E, stage *iii*).

### Fates of luminally released APs

Once released into the lumen, the ER-APs used in this study are stable (t_1/2_ > 8 hr) and are not subject to lysosomal degradation, secretion, or ERAD (Fig 2A, E, F). This is likely due to their being compact structures composed of autonomously folded domains and disordered linkers, and therefore lack ERAD degrons or export motifs. Our data with various reporters demonstrates that the ER-RQC pathway is the primary pathway for degradation of stalled secreted proteins, regardless of the type of luminal cargo. If ER-RQC fails and ER-APs are released into the ER lumen, multiple secondary degradation pathways like ERAD, ER-phagy, or trafficking to the lysosome may target the APs (5, 73, 74). The structural features of the AP may influence its secondary degradation fate, but do not affect its susceptibility to undergo facilitated backsliding.

Unlike the reporters used in this study, endogenous stalls may occur at locations that could disrupt protein secondary structure, thereby generating ERAD degrons. If UFMylation is disrupted, then luminal release of these ERAD degron-containing stalled polypeptides is likely to compete with other ERAD clients. Indeed, we previously identified the entire UFMylation pathway in a CRISPR screen for genes whose disruption stabilizes luminal ERAD reporters (47, 75). The data presented here suggest that UFMylation-dependent, RQC-facilitated backsliding is the primary mechanism for degrading ER-APs, although there are likely redundant ways by which cells can destroy ER-APs once released into the lumen.

### Facilitated backsliding of folded domains

One of the surprising findings of the present work is the observation that ER-APs are dislocated to the cytosol despite the presence of folded domains (VHP, GFP) and core N-glycans. Although SEC61 is permissive for bidirectional nascent chain movement (76), the (∼20-25Å) aperture of this channel is too narrow to permit passive movement of even small folded regions (77, 78). A previous study conducted in cell-free extracts reported that ribosome stalled nascent chains can passively backslide up to - but not beyond - the nearest stably folded VHP domain (4). Our data (Fig 2C), establish that, in intact mammalian cells, this passive backsliding process is supplemented by unfolding forces generated by p97/VCP. Such AAA+ ATPase machines, present in all domains of life, provide sufficient force to completely unfold or extract most proteins (79, 80). p97/VCP engages its substrates via avid binding to polyubiquitin chains on the client (66), so it is likely that AP dislocation through SEC61 must be coupled to polyubiquitylation. While it is possible that, in some cases, a lysine-containing unstructured AP region could extend through the small gap between the translocon and the ribosome to be accessed by LTN1 as proposed (4), our constructs lack unfolded domains of sufficient length to span this gap. Instead, we propose that the strict dependence of our ER-APs on CATylation for degradation suggests that CAT tails facilitate efficient ubiquitylation by LTN1. Considering our finding that ER-AP CAT tails are composed of 15-20 amino acids (Fig S3B-D) and that the average stalled AP contains ∼15 poly(A) encoded lysine codons buried within the ET (11, 19), it is likely that CATylation could push enough lysines out of the ET to encounter the LTN1 RING domain and its cognate E2 (31).

### Essential role of RPL26 UFMylation in ER-RQC

Our data unveil an essential role of RPL26 UFMylation in facilitating ER-RQC. We find that CATylation and UFMylation are essential for and upstream of p97/VCP-dependent ER-AP extraction and UPS-mediated degradation. Our discovery that LTN1 and UFMylation act together to promote ER-AP clearance indicates that UFMylation and RQC operate in the same pathway. Conjugation of UFM1 to RPL26 lysines 132/134 places it immediately adjacent to the physical interface between the ET and the OST/SEC61 translocon complex (5, 47). This position raises the possibility that UFMylation could disrupt the high-affinity interaction between the 60S and the translocon (81, 82), either by the presence of the UFM1 conjugate alone - or more likely given its small size - by recruiting a UFM1-interacting protein (“UFIP” in Fig 5E, *iii*). Disruption of the RTJ could enable bulky RQC and UPS machinery including LTN1 together with its cognate activated E2, or megadalton components such as p97/VCP and the 26S proteasome to access the ER-AP at the RTJ and promote ubiquitin-dependent ATP-driven unfolding of luminal domains, backsliding, and proteolysis. Identification of the putative UFM1-interacting protein and the fine details of the mechanism of RTJ disruption by RPL26 UFMylation merits attention in future studies.

## Materials and Methods

### Mammalian cell culture

HEK293 human embryonic kidney cells (ATCC) and U2OS Human Bone Osteosarcoma Epithelial Cells (ATCC) were maintained in DMEM-High glucose (Cytiva) supplemented with 10% FBS. Cell lines were grown in a humidified incubator at 37°C and 5% CO_2_. All cell lines were routinely tested for mycoplasma infection using a PCR mycoplasma detection kit according to the manufacturer’s instructions (ABM Inc.).

### Mammalian cell transfections

For reporter transfections, HEK293 or U2OS cells were transfected using Lipofectamine 3000 (Thermo Fisher Scientific L3000015) according to the manufacturer’s instructions. Transfected cells were cultured for 24-48 hr before being processed for downstream analysis. Rescue experiments were performed by subjecting HEK293 cells to two rounds of transfection with rescue plasmids for 96 hr. Ribosome stalling reporter transfection was performed for 24-48 hr before being processed for downstream analysis.

### Cell Line Generation

To generate *LTN1^KO^* and *NEMF^KO^* HEK293 cells, we produced sgRNA lentivirus by combining pMCB320 (75) containing the indicated sgRNA with third-generation lentiviral packaging mix (1:1:1 mix of pVSV-G, pMDL, pRSV) and transfected into HEK293T cells using TransIT LT1 transfection reagent (Mirus Bio LLC) according to manufacturer’s protocol. Transfected cells were grown for 72 hr and then the supernatant containing the viral particles was collected and passed through a sterile 0.45 µm Millex syringe filter (Millipore). HEK293 FLAG-Cas9 cells (a gift from Alice Ting, Stanford) were infected by plating cells into complete DMEM with viral supernatant supplemented with 10 µg/mL polybrene and incubated for 72 hr. Infected cells were selected by growing in DMEM containing 2 µg/mL puromycin (Thermo Fisher Scientific) for 3-5 days or until the control, non-transduced cells all died. Cells were recovered in complete DMEM lacking puromycin for 48-72 hr. sgRNA-expressing cell lines were assayed within 10 days or were used to generate clonal cell lines by limited dilution cloning. The clonal cell lines used in this study were screened by immunoblot for loss of the endogenous protein.

To generate the U2OS UBA5^DD^ cells, the *E. coli* dihydrofolate reductase (DHFR) sequence with three destabilizing mutations (R12Y/G67S/Y100I) (70) fused to the N-terminus of *UBA5* was subcloned into pMCB497-pTRE vector (75) using a standard restriction enzyme cloning procedure. U2OS T-Rex Flp-In parental cells (a gift from James Olzmann, UC Berkeley) were transduced with virus containing the pTRE-Tight-DHFR-UBA5, and stable pools were established following selection in complete DMEM supplemented with 7.5 µg/mL Blasticidin for 10 days. Subsequently, knockout of the endogenous *UBA5* gene was performed as described previously (47, 64). U2OS cells expressing doxycycline (DOX)-activated DHFR-UBA5 were cotransfected with two plasmids: (1) pX330-U6-Chimeric BB-CBh-hSpCas9 (Addgene plasmid #42230, a gift from Feng Zhang), for expression of human codon-optimized SpCas9 and sgRNA UBA5 (ACCTACTATTGCTACGGCAA); (2) UBA5 pDONOR-STOP, for homology directed repair (HDR)-mediated insertion of tandem stop codons. DHFR-UBA5 was made to be sgRNA-resistant. Clonal cell lines were selected by limiting dilution after selection with 100 µg/mL hygromycin for 10 days. Expression of the DHFR-UBA5 fusion protein was sustained by growing cells in the presence of 1 mg/mL DOX and the small-molecule ligand trimethoprim (TMP) (10 µM), which stabilizes the destabilized DHFR domain, to ensure that U2OS UBA5^DD^ cells were never exposed to UBA5 depletion. Single cell-derived clones were expanded under selective pressure and screened by immunoblotting analysis of UBA5. In the washout experiments, cells were gently rinsed three times in complete media to remove the TMP ligand and DOX to rapidly degrade DHFR-UBA5, and incubated in complete DMEM lacking TMP and DOX for up to 24 hr (i.e. “+Washout condition”). Other cell lines used in this study are listed in the Key Resources Table.

### Plasmids

To generate stalling reporter sequences, FLAG-VHP-sequon-VHP-K20-GFP-HA was ordered as a gene block (Genscript) and inserted into a pcDNA3.1 parent vector containing a CMV promoter. The signal sequence of bovine preprolactin and the K0 sequence were purchased as oligos which were annealed and subcloned into the reporter plasmid upstream of the FLAG tag or downstream of the VHP domains, respectively. The N-glycosylation sites (sequons) were added or removed by site directed mutagenesis. The other variants of the reporters were generated by standard subcloning methods. The original reporter design generated a frameshift (FS) species which migrated close to the AP species (MW: ∼25kDa) (Original reporter used in Fig 2E, S2A). Mutagenesis was performed to remove out of frame stop codons from GFP, making the FS species a greater molecular weight, which runs slower than the RT product, named “FS-Corrected (MW: ∼60kDa) (FS-Corrected reporters are used in all other experiments, see labeled FS-Corrected species in Fig S2F). To generate SS^GFP^, EGFP and the spacer was amplified from the ER-K20 reporter used in Wang et al, 2020 (5) and inserted into SS^VV^-K20 after the preprolactin signal sequence and before the K20 stall sequence.

The NEMF rescue plasmids, NEMF-WT and NEMF-DR (a gift from Claudio Joazeiro, Heidelberg University) were mutated with synonymous mutations to become sgResistant.

### Glycosidase treatment

HEK293 cells were collected and lysed in RIPA lysis buffer (50 mM Tris pH 7.6, 150 mM NaCl, 1% NP-40, 0.5% Sodium Deoxycholate, 0.1% SDS) with protease inhibitor cocktail and 1mM PMSF. Total protein concentration was determined for each sample with the Pierce BCA Protein Assay Kit (23225) according to manufacturer’s instructions. Protein concentrations were normalized between samples. Samples were denatured and endoH treated following the manufacturer’s protocols (New England Biolabs, inc. P0702L). Reactions were incubated at 37°C for 1 hr and then analyzed by SDS-PAGE and immunoblotting.

### Translation shut-off assay

HEK293 cells were treated with emetine at a final concentration of 20 µM for the indicated times. For treatment with Bortezomib (BTZ), 1 µM BTZ was added simultaneously with emetine. Cells were collected and washed in PBS after which they were lysed in RIPA lysis buffer (50 mM Tris pH 7.6, 150 mM NaCl, 1% NP-40, 0.5% Sodium Deoxycholate, 0.1% SDS) with protease inhibitor cocktail and 1mM PMSF. Total protein concentration was determined for each sample with the Pierce BCA Protein Assay Kit (23225) according to manufacturer’s instructions. Equal amounts of total protein were analyzed by immunoblotting.

### Sucrose cushion sedimentation

Cells were collected and lysed in 1% Triton lysis buffer (20 mM Tris pH 7.5, 150 mM NaCl, 5 mM MgCl_2_, 1% Triton X-100) with protease inhibitor cocktail, 1 mM PMSF, and 1 mM DTT. Total protein concentration was determined for each sample with the Pierce 600 nm Protein Assay Reagent according to the manufacturer’s protocol. Samples were centrifuged at 100,000 × g for 1 hr at 4°C through a 1 M sucrose cushion in 1% Triton lysis buffer. Pellets were washed once with ice cold H_2_O and resuspended in 1X Laemmli buffer containing 2-mercaptoethanol 5% (v/v) by heating at 100°C for 5 min. Samples were analyzed by SDS-PAGE and immunoblotting.

### Cell fractionation by sequential detergent extraction

U2OS or HEK293 cell fractionation was performed as described previously (47) with some modifications. Cells were collected in PBS and centrifuged at 800 × g for 5 min and cell pellets (∼2 × 10^6^) were resuspended in 150 µL of “fractionation” buffer (50 mM Hepes pH 7.35, 150 mM NaCl) with protease inhibitor cocktail and 1 mM PMSF. One third of the sample was lysed in a fractionation buffer containing 1% NP-40 for 10 min on ice, followed by centrifugation for 10 min at 21,130 at 4°C to obtain a clear whole cell lysate sample (“WCL”). The remaining two thirds of the sample were resuspended in 100 µL of fractionation buffer containing 0.005% digitonin for 5 min to permeabilize the plasma membrane, followed by centrifugation at 8,000 g for 5 min. The supernatant containing the cytosolic fraction was collected (“Cyto”) and the pellet was washed once with 500 µL of fractionation buffer with no detergents before resuspending in 100 µL fractionation buffer containing 1% NP-40. After incubating for 10 min on ice and centrifugation at 21,130 × g for 10 min, the supernatant containing the membrane fraction (“ER”) was collected. In order to separate the ER lumen from the ER membrane we adapted the protocol described above by using a higher concentration of digitonin (i.e., 0.2% rather than 0.005%) to solubilize the luminal content of the ER without extracting ER membrane proteins. Equal volumes of collected fractions were analyzed by SDS-PAGE and immunoblotted.

### SDS-PAGE and immunoblotting

Proteins were denatured in 1X Laemmli buffer containing 2-mercaptoethanol 5% (v/v) by heating at 100°C for 5 min. Samples were then separated by SDS-PAGE (12% Tris-Glycine gels or “4-20% Mini-PROTEAN TGX” (Bio-Rad)) and transferred in a semi-dry transfer to nitrocellulose following manufacturer’s protocol (Bio-Rad). Nitrocellulose membranes were blocked in Intercept® (TBS) Blocking Buffer to reduce nonspecific antibody binding and incubated with primary antibodies diluted in PBS-T containing 0.1% Tween-20 and 5% BSA. Immunoreactivity was detected using fluorescent IRDye secondary antibodies and scanning by Odyssey imaging (Li-COR Biosciences). Band intensities were quantified by Image Studio Lite software (Li-COR Biosciences). For translational shut-off assays, bands were quantified by densitometry and protein levels at each time point were normalized to Tubulin or GAPDH. Percentage protein remaining was calculated relative to t = 0 hr for each cell line. Ubiquitylation of RPS10 was assayed as in (15).

### Small interfering RNA (siRNA) knockdown

Lipofectamine RNAiMAX from Thermo Fisher Scientific (13778030) was used for transient siRNA transfections according to manufacturer’s instructions for reverse transfection of RNAi. Silencer Select siRNAs were purchased from Thermo Fisher Scientific: ZNF598 s40510, ASCC3 s21603, SYVN1 (HRD1) s39020, LTN1 s25003, NEMF s17483, and scrambled (negative control no. 1). Cells were analyzed 48-72 hr post-transfection.

### Statistical analysis

Data are represented as the mean ± SD unless otherwise stated. The number of independent replicates performed for each experiment is indicated in the figure legends. Western blot band intensities were quantified using Image Studio Lite version 5.2.5 (LI-COR Biosciences) and normalized to loading control. Protein remaining was calculated as a percentage of time 0 and one-phase exponential decay curves were fit using Prism 9 (GraphPad Software).

## Supporting information

Supplemental data for Scavone et al

## Acknowledgements

We thank Celeste Riepe (Stanford University) and Onn Brandman (Stanford University) for critical reading of the manuscript. We thank Danish Khan (Stanford University), Carolyn Bertozzi (Stanford University), Daniel Hebert (University of Massachusetts), Rachel Green (Johns Hopkins University School of Medicine), Niladri Sinha (Johns Hopkins University School of Medicine), and Anne Bertolotti (MRC laboratory, Cambridge) for helpful discussions. We thank James Olzmann (UC Berkeley) for U2OS T-Rex Flp-In cells, Alice Ting (Stanford University) for FLAG-Cas9 HEK293 cells, Yihong Ye (NIH, Bethesda) for HEK-T Parental and RPL26ΔC cells, Claudio Joaziero (Heidelberg University) for NEMF plasmids, and Ramanujan Hegde (MRC laboratory, Cambridge) for anti-SEC61β antibody. This work was supported by the NIGMS (R01GM074874) to R.R.K., by the NIH (T32GM007276) to S.C.G., and by the Stanford Graduate Fellowship to S.C.G.

