## Supplemental data for Scavone et al for "RPL26/uL24 UFMylation is essential for ribosome-associated quality control at the endoplasmic reticulum"

**SI Appendix:**

#### Table of Contents

Figure Legends S1 to S4
Figures S1 to S4
Table of Reagents

**Figure S1. Related to Figure 1. Characterization of ER-targeted ribosome stalling reporters.**

**A:** The UFMylation system shown in relation to the ER membrane bilayer. UFM1 conjugation machinery (E1: UBA5, E2: UFC1, E3: UFL1, CDK5RAP3, DDRGK1) and deconjugation machinery (UFSP1, UFSP2, ODR4) are shown in their associated complexes. The target of UFMylation, RPL26, is shown in the context of the 60S ribosomal subunit.

**B:** PNGase F treatment on $SS^{gVgV}$ reporter confirms correct ER targeting of $SS^{gVgV}$ by increase in AP and RT mobility. Glycosylated (+2g) and nonglycosylated RT and AP species are indicated; data shown are representative of three independent experiments.

**C:** ER-stalled ribosomes are recognized and split by ZNF598 and ASCC3, respectively. HEK293 cells were transfected with scrambled (SCR), ZNF598, or ASCC3 small interfering RNAs (siRNA) and stalling reporter constructs $Cyto^{VV}$, $SS^{VV}$, $SS^{VgV}$. Immunoblot analysis was performed on WCLs using FLAG antibody to monitor changes in steady-state levels of frameshift (FS), RT, and AP species. Asterisks indicate nonspecific immunoreactive bands. Data shown are representative of three independent experiments.

**D:** Cell fractionation analysis of subcellular $Cyto^{VV}$-AP distribution. HEK293 cells were transfected with $Cyto^{VV}$ and subjected to cell fractionation. Reporter products were analyzed by immunoblot of WCL, Cyto, and ER cell fractions with FLAG antibody. GAPDH: cytosol marker; SEC61β: ER marker; data shown are representative of three independent experiments.

**E:** Titration of digitonin to determine the optimal concentration to solubilize the luminal contents of the ER (Sup) without extracting ER membrane proteins (Pellet). PDI: ER lumen marker; SEC61β and HRD1: ER membrane markers. Red box indicates the optimal digitonin concentration to separate ER luminal proteins (“ER-Lum”) from ER membrane proteins (“ER-Mem”) for the experiment in Figure 1G; data shown are representative of two independent experiments.

**F:** Ribosome association of ER-APs assessed by sedimentation. HEK293 cells were transfected with $SS^{gVgV}$ and subjected to sucrose cushion sedimentation. Reporter products were analyzed by immunoblot of WCL, ribosome-free (Sup), and ribosome-associated (Pellet) fractions with FLAG antibody. RPL17: loading control.

**Figure S2. Related to Figure 2. Validation data for ER-AP degradation via the proteasome.**

**A:** Prolonged BafA treatment does not result in ER-AP stabilization. HEK293 cells were transfected with the ER targeted stalling reporters, $SS^{VgV}$ (“Original” reporter, see figure S2F) and $SS^{VV}$, incubated with DMSO, 1 µM BTZ for 4 hr, or 100 nM BafA for 4 hr or 16 hr. WCLs were analyzed by immunoblot with FLAG antibody to detect AP and AP+g species, and LC3 antibody to assess the effect of BafA treatment on LC3-II

accumulation. GAPDH: loading control; data shown are representative of two independent experiments.
**B:** BTZ treatment promotes accumulation of a nonglycosylated $SS^{VgV}$ ER-AP species. HEK293 cells were transfected with the indicated reporter and treated with DMSO or 1 µM BTZ for 4 hr. Reporter products were analyzed by immunoblot of WCLs with FLAG antibody. Labels indicate mobilities of glycosylated (+g) and non-glycosylated RT and APs; data shown are representative of three independent experiments.
**C:** Cell fractionation analysis of subcellular $Cyto^{VV}$-AP distribution following BTZ treatment. U2OS cells were transfected with the indicated reporter and treated with DMSO or 1 µM BTZ for 4 hr prior to cell fractionation. Reporter products were analyzed by immunoblot of WCL, Cyto, and ER cell fractions with FLAG antibody. GAPDH: cytosol marker; SEC61β: ER marker.
**D:** Validation of BTZ and NMS-873 efficacy. HEK293 cells transfected with the indicated reporters were treated for 4 hr with 1 µM BTZ or 5 µM NMS-873. Reporter products were analyzed by immunoblot with FLAG antibody. Immunoblotting with anti-CD147 antibody was performed to assess the effect of BTZ and NMS-873 on steady state levels of CD147, an endogenous proteasome substrate. Mat: Mature glycosylated CD147; CG: Core Glycosylated CD147; Degly: Deglycosylated CD147; data shown are representative of two independent experiments.
**E**: A GFP-based stalling reporter is degraded by the proteasome. *Upper panel,* Schematic of GFP-based stalling reporter, $SS^{GFP}$. Reporter contains $SS^{PPL}$, signal sequence from bovine preprolactin; EGFP, enhanced green fluorescent protein; FLAG, FLAG epitope tag. Composition of the reporter is shown below. The predicted MW for arrest peptide (AP, black line) is 50 kDa. *Lower panel,* HEK293 cells expressing $SS^{GFP}$ were treated either with DMSO, 1 µM bortezomib (BTZ), or 100 nM Bafilomycin A1 (BafA) for 4 hr. WCLs were analyzed by immunoblot with FLAG antibody to detect AP species, and LC3 antibody to assess the effect of BafA treatment on LC3-II accumulation. GAPDH: loading control. *Right Panel,* Topological organization of GFP arrest peptide when stalled at the translocon. When the ribosome translating $SS^{GFP}$ reaches the stall sequence, the entire GFP protein will be translocated into the ER lumen and folded.
**F:** Schematic of the two reporter variants illustrating the frameshift (FS) species generated by our stalling reporters as described in the materials and methods section. Original: frameshift product generated by this reporter is ~25kD. FS-corrected: frameshift product generated by this reporter is 60kD or 65kD. S Tag+1 and S Tag+2 are generated by out of frame translation downstream of the stalling sequence (K20). The Original reporter was used in Figures 2E, S2A. The FS-corrected reporter is used in all other experiments.

**Figure S3. Related to Figure 3. Extended data for the requirement of RQC machinery to degrade ER-APs.**

**A:** Control for Fig 3A. Effect of scrambled (SCR), LTN1, or HRD1 siRNA on endogenous protein levels, assayed by immunoblot for endogenous LTN1 or HRD1 with anti-LTN1 or anti-HRD1 antibody, respectively.

**B-D:** Estimation of CAT tail length for ER-APs.

**B:** Representative 12% SDS-PAGE used to determine the molecular weight (MW) of ER-AP CAT tails. As in Fig 3E, HEK293 $NEMF^{KO}$ cells were rescued with WT NEMF or NEMF-DR and transfected with the $SS^{VV}$ stalling reporter. WCLs were analyzed by immunoblot with anti-FLAG antibody. Unmodified APs are indicated by the label “AP” and by black arrowheads; CATylated APs are indicated by “$AP^{CAT}$”. Blue arrows indicate migration distance (inches) from the top of the gel. GAPDH and tubulin: loading controls.

**C:** Semi-logarithmic plot of MW of size markers versus migration distance (inches) from Fig S3B. Data shown are representative of four experiments. The equation of the best fit line (y = -0.34x + 2.03) is used to calculate the MW of the AP species in this experiment.

**D:** Summary table of four independent measurements for $SS^{VV}$ CAT tail length. MW of AP or $AP^{CAT}$ was estimated by linear regression analysis of semi-logarithmic plots of MW of size markers versus migration distance of AP or $AP^{CAT}$ in 12% SDS-PAGE.

**Figure S4. Related to Figure 4. Extended data on the requirement of UFMylation for ER-AP degradation.**

**A:** ER-targeted stalls specifically promote RPL26 UFMylation. *Upper panel,* HEK293 cells were transfected with 2 µgs of empty vector (EV) or indicated nonstall (K0) or stall (K20) reporter plasmids. 20% of WCLs from samples transfected with K0 reporters were loaded compared to WCLs transfected with K20 stall reporters. Where indicated cells were treated with 200 nM ANS (+ANS) for 15 minutes prior to cell lysis. WCLs were analyzed by immunoblot with anti-FLAG antibody. *Lower panel,* WCLs were sedimented through a sucrose cushion and the pellets were analyzed by immunoblot with anti-UFM1 antibody. RPL17: loading control.

**B:** Cytosolic stalling reporter does not induce UFMylation. *Upper panel,* HEK293 cells were transfected with 4 or 5 µgs of empty vector (EV) or $Cyto^{VV}$-K20. Where indicated cells were treated with 200 nM ANS (+ANS) for 15 minutes prior to cell lysis. GAPDH: loading control. *Lower panel,* WCLs were sedimented through a sucrose cushion and the pellets were analyzed by immunoblot with anti-UFM1 antibody. RPL17: loading control.

**C:** ER stalling reporters with diverse sequences and topologies induce RPL26 UFMylation. *Left panels,* Schematic of the stalling reporters used in this experiment. $TFR^{TMD}$, transmembrane domain from transferrin receptor; $SS^{PPL}$, signal sequence from bovine preprolactin; $PPL^{FL}$, prolactin; FLAG, FLAG epitope tag; glyc, position of an N-glycosylation site or “sequon”; “Y”, N-glycan; VHP, villin headpiece domain; K20, indication that the reporter contains a polylysine stalling sequence of 20 lysine residues; GFP, superfolder green fluorescent protein; V5, epitope tag; HA, epitope tag. Composition of each reporter shown below, with predicted MW for arrest peptide (AP, black line) or readthrough (RT, black line + dashed black line) species produced by each stalling reporter. *Right panel,* HEK293 cells were transfected with 2µgs of empty vector (EV) or the indicated stall (K20) reporters. Where indicated cells were treated with 200 nM ANS (+ANS) for 15 minutes prior to cell lysis. WCLs were analyzed by immunoblot with anti-FLAG antibody. Tubulin: loading control. *Lower panel,* WCLs were sedimented through a sucrose cushion and the pellets were analyzed by immunoblot with anti-UFM1 antibody. RPL17: loading control.

**D:** $UFM1^{KO}$ stabilizes ER but not cytosolic-APs in U2OS cells, as in HEK293 cells (figure 4D). U2OS WT and clonal $UFM1^{KO}$ cell lines were transfected with the indicated reporters. Reporter products were analyzed by immunoblot with FLAG antibody. Knockout was confirmed by immunoblot with anti-UFM1 antibody. GAPDH: loading control.

**E:** ER-AP degradation requires UFM1 conjugation machinery. HEK293 WT, $UFM1^{KO}$, $UFC1^{KO}$, and $UFL1^{KO}$ cell lines were transfected with the indicated reporters. Reporter products were analyzed by immunoblot with FLAG antibody. Knockouts were confirmed by blotting with antibodies against endogenous UFC1 or UFL1 proteins with anti-UFC1 or anti-UFL1 antibodies, respectively. GAPDH: loading control; data shown are representative of two independent experiments.

**F:** Validation of U2OS $UBA5^{DD}$ cells. UBA5 and RPL26-UFM1 are undetectable 24 hr after washout. U2OS $UBA5^{DD}$ cells were cultured in complete DMEM with TMP ("-Washout") or washed to remove TMP from the media ("+Washout") at the indicated times. *Upper panel,* WCLs were sedimented through a 1M sucrose cushion and the pellets were analyzed by immunoblot with anti-UFM1 antibody. RPL17: loading control. *Middle panel,* WCLs from this experiment were analyzed by immunoblot with anti-UBA5 antibody to assess levels of remaining DHFR-UBA5. *Lower panel,* WCLs were analyzed by immunoblot with anti-UFM1 to detect conjugates with UFC1 and DHFR-UBA5 which are lost after acute depletion of UBA5. Asterisks indicate nonspecific immunoreactive bands; data shown are representative of two independent experiments.

### Supplemental Figure 1

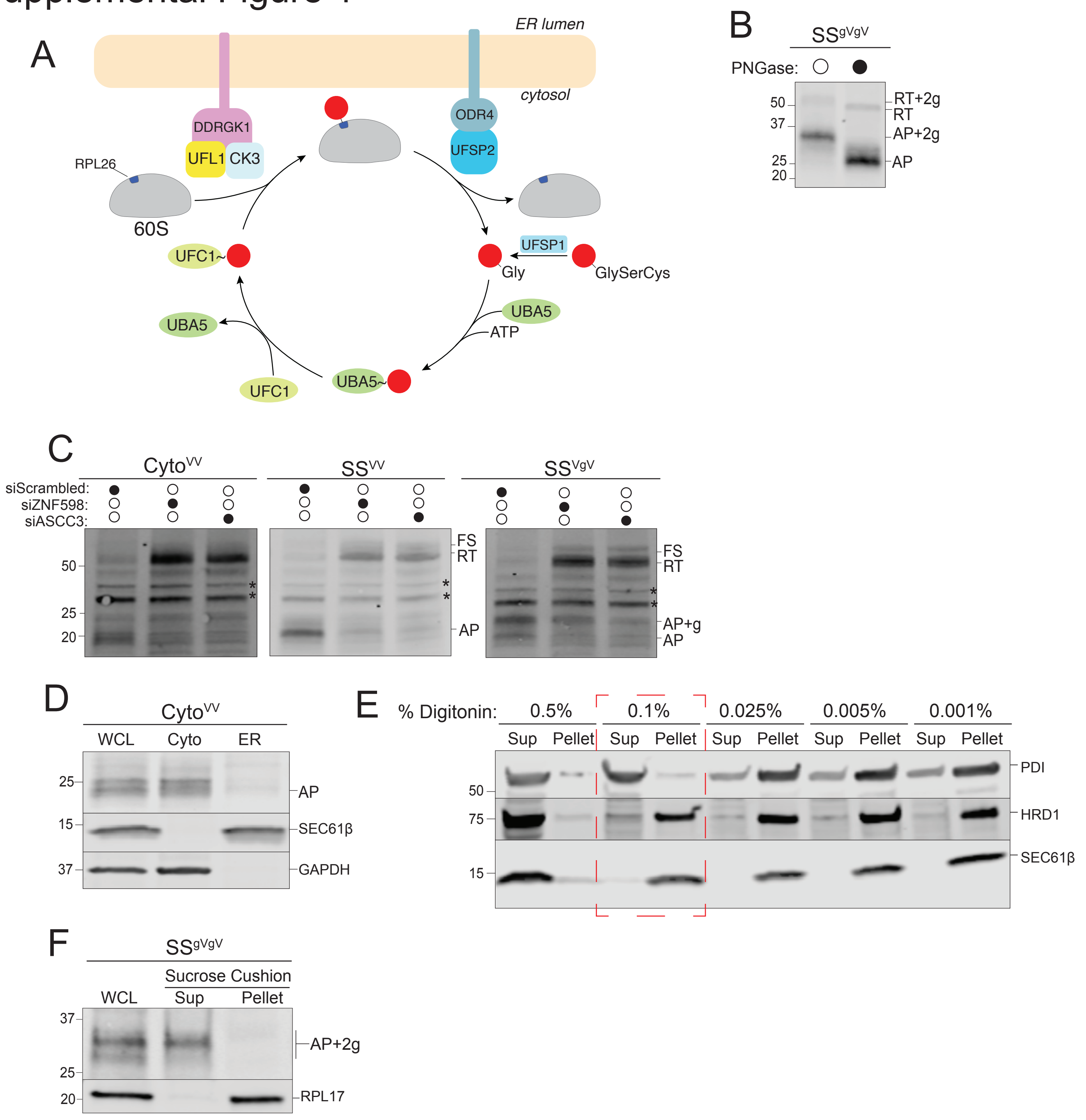

### Supplemental Figure 2

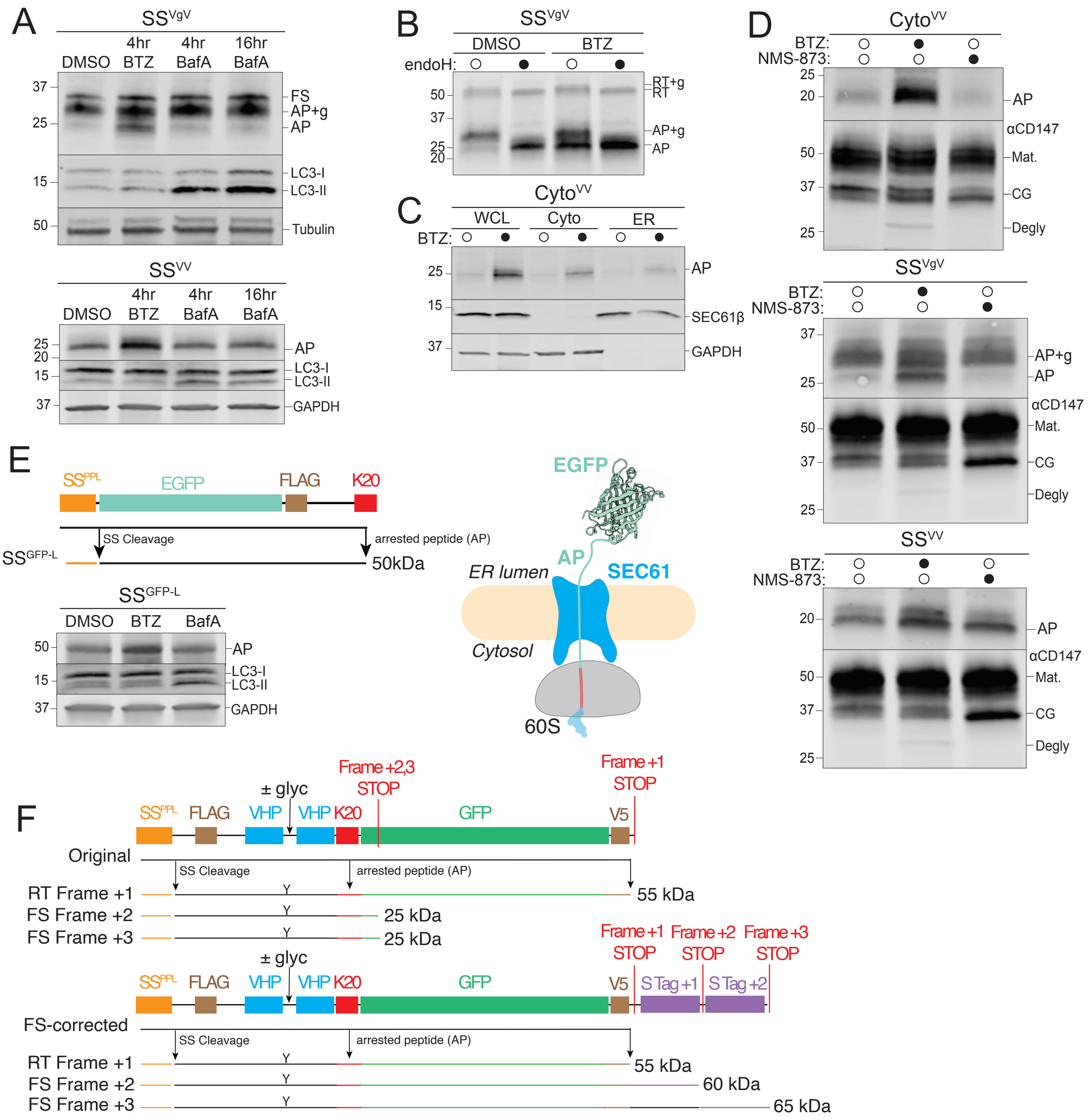

### Supplemental Figure 3

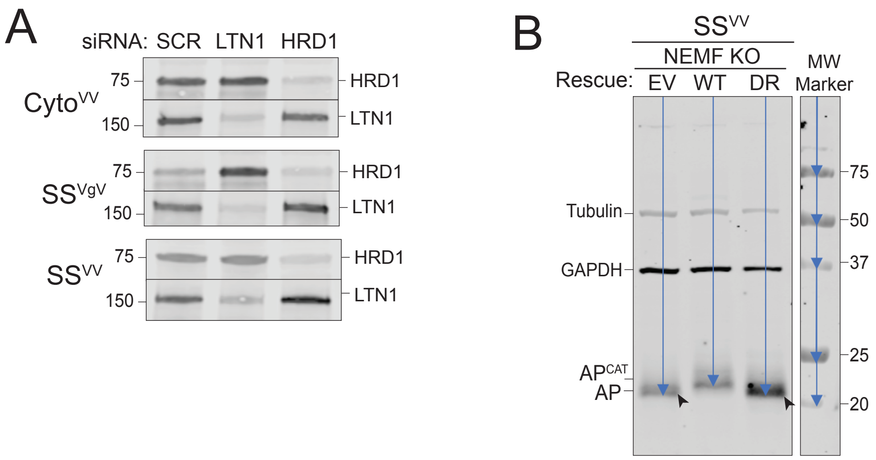

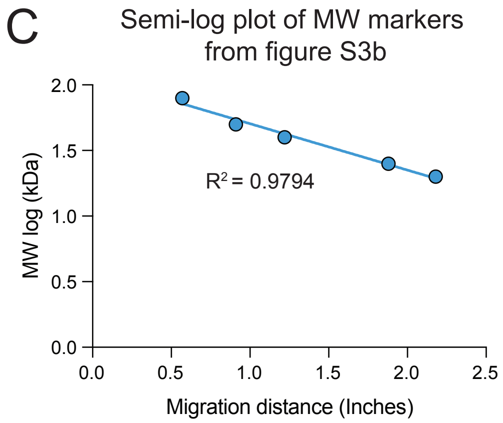

D

| | Expt 1 | Expt 2 | Expt 3 | Expt 4 |
|---|---|---|---|---|
| $AP^{CAT}$(kDa) | 22.2 | 21.7 | 21.4 | 23.4 |
| AP(kDa) | 20.9 | 20.5 | 20.2 | 22.1 |
| CAT tail length (kDa) | 1.3 | 1.2 | 1.2 | 1.3 |

### Supplemental Figure 4

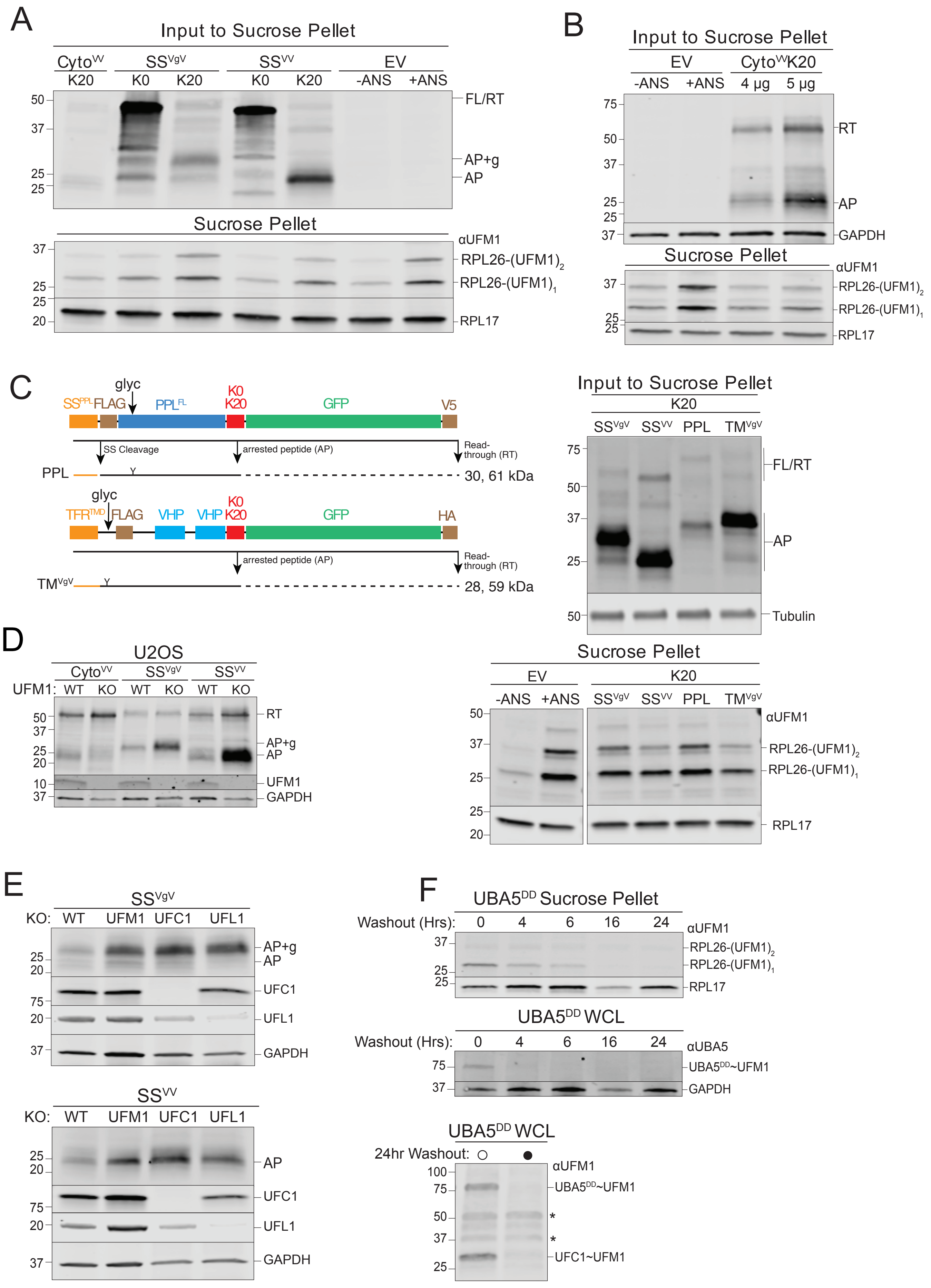

| Reagent or Resource | Source | Identifier (CAT #) |
|---|---|---|
| Antibodies | | |
| Mouse monoclonal anti-FLAG | Sigma-Aldrich | F1804 |
| Mouse monoclonal anti-GAPDH | Cell Signaling | 97166S |
| Rabbit polyclonal anti-GAPDH | Cell Signaling | 2118S |
| Mouse monoclonal anti-PDI | Enzo Life Sciences | ADI-SPA-891 |
| Rabbit polyclonal anti-SEC61β | Gift from Hegde Lab | N/A |
| Rabbit polyclonal anti-RPS10 | Abclonal technologies | A6056 |
| Rabbit polyclonal anti-ASCC3 | Proteintech | 17627-1-AP |
| Rabbit polyclonal anti-Tubulin | Abcam | ab15246 |
| Rabbit polyclonal anti-HRD1 | Proteintech | 13473-1-AP |
| Mouse monoclonal anti-RPL17 | Santa Cruz Biotechnology | sc-515904 |
| Rabbit polyclonal anti-LC3B | Cell Signaling | 2775S |
| Rabbit polyclonal anti-SEL1L | [illegible] Abcepta) | N/A |
| Mouse monoclonal anti-CD147 | Santa Cruz Biotechnology | sc-21746 |
| Rabbit polyclonal anti-LTN1 | Proteintech | 28452-1-AP |
| Rabbit polyclonal anti-NEMF | Thermo Fisher Scientific | PA5-36308 |
| Rabbit monoclonal anti-UFM1 | Abcam | ab109305 |
| Rabbit monoclonal anti-UFC1 | Abcam | ab189252 |
| Rabbit monoclonal anti-UFL1 | Abcam | ab227506 |
| Rabbit polyclonal anti-UBA5 | Proteintech | 12093-1-AP |
| Rabbit polyclonal anti-RPL26 | Abcam | ab59567 |
| Goat anti-Mouse IgG, IRDye 800CW | LI-COR Biosciences | 926-32210 |
| Goat anti-Mouse IgG, IRDye 680LT | LI-COR Biosciences | 926-68020 |
| Goat anti-Rabbit IgG, IRDye 800CW | LI-COR Biosciences | 926-32211 |
| Goat anti-Rabbit IgG, IRDye 680LT | LI-COR Biosciences | 926-68021 |
| Bacterial and Virus Strains | | |
| E. coli: MAX efficiency DH5α competent cells | Thermo Fisher Scientific | 18258012E |
| Chemicals, Peptides, and Recombinant Proteins | | |
| cOmplete, EDTA-free Protease Inhibitor Cocktail | Sigma-Aldrich | 11873580001 |
| Doxycycline hyclate | Sigma-Aldrich | D9891 |
| Lipofectamine 3000 | Thermo Fisher Scientific | L3000015 |
| Lipofectamine RNAiMAX | Life Technologies/Invitrogen | 13778100 |
| Emetine dihydrochloride hydrate | Sigma-Aldrich | E2375 |
| NMS-873 | Sigma-Aldrich | SML1128 |
| Puromycin dihydrochloride | Thermo Fisher Scientific | A1113803 |
| Bortezomib | Selleck Chemicals | S1013 |
| Phenylmethylsulfonyl fluoride (PMSF) | Sigma-Aldrich | 10837091001 |
| Odessey Blocking Buffer | LI-COR Biosciences | 927-60003 |
| Endo H | New England Biolabs | P0702S |
| PNGase F | New England Biolabs | P0704S |
| N-ethylmaleimide | Sigma-Aldrich | E3876 |
| Phusion Master Mix with HF Buffer | Thermo Fisher Scientific | F-531L |
| Dithiothreitol (DTT) | Fisher Scientific | ab000490-00010 |

| | | |
|---|---|---|
| TransIT LT1 transfection reagent | Mirus Bio LLC | MIR 2300 |
| Anisomycin | Sigma-Aldrich | A9789 |
| Bafilomycin A1 | Sigma-Aldrich | B1793 |
| Digitonin | EMD Millipore | 300410 |
| Sucrose | Sigma-Aldrich | S8501 |
| TMP | Sigma-Aldrich | T7883 |
| HyClone Dulbecco's Modified Eagle Medium (DMEM) with high glucose | Cytiva | SH30285.FS |
| Critical Commercial Assays | | |
| PCR Mycoplasma Detection Kit | ABM inc. | G238 |
| BCA Protein Assay Kit | Thermo Fisher Scientific | 23225 |
| 660 nm Protein Assay Reagent | Thermo Fisher Scientific | 22660 |
| QIAprep Spin Miniprep Kit | Qiagen | 27106X4 |
| PureLink™ HiPure Plasmid Midiprep Kit | Thermo Fisher Scientific | K210004 |
| Experimental Models: Cell Lines | | |
| Human: HEK293 Cells | ATCC | CRL-1573 |
| Human: HEK293T Cells | ATCC | CRL-3216 |
| Human: U2OS Cells | ATCC | HTB-96 |
| Human: HEK293 HRD1 KO Cells | (van der Goot et al., 2018) | N/A |
| Human: HEK293 OS9 KO Cells | (van der Goot et al., 2018) | N/A |
| Human: HEK293 SEL1L KO Cells | (van der Goot et al., 2018) | N/A |
| Human: HEK293 LTN1 KO Cells | This study | N/A |
| Human: HEK293 NEMF KO Cells | This study | N/A |
| Human: HEK293 UFM1 KO Cells | (Walczak et al., 2019) | N/A |
| Human: HEK293 UFC1 KO Cells | (Walczak et al., 2019) | N/A |
| Human: HEK293 UFL1 KO Cells | (Walczak et al., 2019) | N/A |
| Human: U2OS UBA5-DD Cells | This study | N/A |
| Human: U2OS UFM1 KO Cells | (Walczak et al., 2019) | N/A |
| Human: HEK293 FLAG-Cas9 Cells | Gift from Ting Lab | N/A |
| Human: HEKT Cells | Gift from Yi Lab | N/A |
| Human: HEKT RPL26ΔC | Gift from Yi Lab | N/A |
| Oligonucleotides | | |
| Silencer™ Select Negative Control No. 1 siRNA | Thermo Fisher Scientific | 4390843 |
| siRNA against ZNF598 | Thermo Fisher Scientific | 4392420 |
| siRNA against ASCC3 | Thermo Fisher Scientific | s21603 |
| siRNA against LTN1 | Thermo Fisher Scientific | s25003 |
| siRNA against HRD1 | Thermo Fisher Scientific | 4427037 |
| siRNA against NEMF | Thermo Fisher Scientific | s17485 |
| SS Bovine Preprolactin Fw:<br>GGATCCACCatggacagcaaaggttcgtcgcagaaagggtcccgcctgctcctgctgctggtggtgtcaaatctactcttgtgccagggtgtggtctccacc<br>TCAGGGTCTGGTAGCGGCGGCCGC | This study | N/A |
| SS Bovine Preprolactin Rev:<br>GCGGCCGCCGCTACCAGACCCTGAggtggagaccacaccctggcacaagagtagatttgacaccaccagcagcaggagcaggcgggaccc<br>tttctgcgacgaacctttgctgtccatGGTGGATCC | This study | N/A |
| K20 Stall Sequence Fw:<br>CAAAAAAAAAAAAAAAAAAAAAAAAAAAAAAAAAAAAAAAAAAAAAAAAAAAAAAAAAAAAAGCTAAGCC | This study | N/A |
| K20 Stall Sequence Rev:<br>GGCTTAGCTTTTTTTTTTTTTTTTTTTTTTTTTTTTTTTTTTTTTTTTTTTTTTTTTTTTTTTTTG | This study | N/A |

| | | |
|---|---|---|
| Recombinant DNA | | |
| sgRNA NEMF (sequence: GACCTCCGCGCCGTACTCG) | This study | N/A |
| sgRNA LTN1 (sequence: AATGCCGAACGACAGTGAA) | This study | N/A |
| sgRNA UBA5 (sequence: ACCTACTATTGCTACGGCAA) | (Walczak et al., 2019) | N/A |
| Plasmid: pcDNA3.1 (+) | Thermo Fisher Scientific | V79020 |
| Plasmid: UBA5 pDONOR-STOP | (Walczak et al., 2019) | N/A |
| Plasmid: ecDHFR(R12Y, G67S, Y100I)-UBA5(sgResistant) | This study | N/A |
| Plasmid: NEMF WT (sgResistant) | (Thrun et al., 2021) | N/A |
| Plasmid: NEMF DR mutant (sgResistant) | (Thrun et al., 2021) | N/A |
| Plasmid: Cyto-VV-K0 | This study | N/A |
| Plasmid: Cyto-VV-K20 | This study | N/A |
| Plasmid: SS-VV-K0 | This study | N/A |
| Plasmid: SS-VV-K20 | This study | N/A |
| Plasmid: SS-VgV-K0 | This study | N/A |
| Plasmid: SS-VgV-K20 | This study | N/A |
| Plasmid: SS-gVgV-K20 | This study | N/A |
| Plasmid: TM-VgV-K0 | This study | N/A |
| Plasmid: TM-VgV-K20 | This study | N/A |
| Plasmid: PPL-K0 | This study | N/A |
| Plasmid: PPL-K20 | This study | N/A |
| Plasmid: SS-GFP-L | This study | N/A |
| Plasmid: SS-GFP-S | This study | N/A |
| Software and Algorithms | | |
| Image Studio Lite version 5.2.5 | LI-COR Biosciences | https://www.licor.com/bio/image-studio/ |
| Prism 9 | GraphPad | https://www.graphpad.com/scientific-software/prism/ |
| SnapGene | Dotmatics | https://www.snapgene.com/ |
| Other | | |
| Trans-Blot Turbo RTA Midi NitrocelluloseTransfer Kit | Bio-Rad Laboratories | 1704271 |
| 4-20% Mini-PROTEAN TGX Gel 10 well | Bio-Rad Laboratories | 4561094 |
| 4-20% Mini-PROTEAN TGX Gel 15 well | Bio-Rad Laboratories | 4561096 |
